# Coevolution of *Drosophila*-type Timeless with Partner Clock Proteins

**DOI:** 10.1101/2024.12.25.628932

**Authors:** Enrico Bullo, Ping Chen, Ivan Fiala, Vlastimil Smýkal, David Doležel

## Abstract

*Drosophila*-type timeless (dTIM) is established key clock protein in fruit flies, regulating the rhythmicity and light-mediated entrainment. However, as indicated by functional experiments, its contribution to the clock differs in various insects. Therefore, we conducted a comprehensive phylogenetic analysis of dTIM across animals, dated its origin, gene duplications, and losses. We identified variable and conserved protein domains, and pinpointed animal lineages that underwent the biggest changes in the dTIM sequence. While dTIM modifications are only mildly affected by changes in the PER protein, even the complete loss of PER in echinoderms had no impact on dTIM. However, changes in dTIM always co-occur with the loss of CRYPTOCHROMES or JETLAG. This is exemplified by the remarkably accelerated evolution of dTIM in phylloxera and aphids. Finally, alternative *d-tim* splicing, characteristic of *D. melanogaster* temperature-dependent function, is conserved at least to some extent in Diptera, albeit with unique alterations. Altogether, this study pinpoints major changes that shaped dTIM origin and evolution.

## INTRODUCTION

The majority of life forms are exposed to periodic alternation of day and night. Anticipating environmental changes such as dawn, dusk, or midday heat provides a significant advantage to various organisms. Circadian clocks, the molecular devices ‘ticking’ with an approximately 24-hour period, have evolved independently in cyanobacteria, plants, fungi, and animals (Bhadra et al., 2017; Dunlap 1999). Despite their independent origins, some molecular mechanisms appear to be conserved (Edgar et al., 2012). The animal circadian clock, best characterized in the fruit fly *Drosophila melanogaster* and mouse *Mus musculus*, operates through several interconnected transcription-translation feedback loops (TTFL) (Dunlap 1999; Mendoza-Viveros et al. 2016). These clock components are broadly conserved across bilaterian animals, with some components traceable back to cnidarians (Kwiatkowski and Emery, 2024; Aguillon et al., 2024. Key transcription factors, such as CLOCK and CYCLE (with the transactivation domain of CYCLE uniquely lost in *Drosophila* and other cyclorrhaphan flies) or CLOCK and BMAL1 (in most other species), are integral to the circadian clock (Thakkar et al., 2022; Tomioka and Matsumoto, 2015). These transcription factors belong to the basic helix-loop-helix (bHLH) Period- Arnt-Singleminded (PAS) protein family and act as critical activators of the clock (Rutila et al., 1998; Allada et al., 1998; Michael et al., 2023; Tumova et al., 2024).

CLK and CYC/BMAL1 drive the expression of both positive and negative regulators in the circadian clock. In *Drosophila*, the key negative regulators are the PERIOD (PER) and *Drosophila*-type TIMELESS (dTIM) proteins. In contrast, the mammalian clock relies on PER proteins (three paralogs in mice) and mammalian-type CRYPTOCHROMEs (mCRYs; two paralogs in mice), sometimes referred to as CRY2-type (Kume et al. 1999; Putker et al. 2021; Cai and Chiu 2021). CRYs are phylogenetically related to photolyases, although mCRYs have lost their ability to detect light (Yuan et al. 2007). The *Drosophila*-type CRY (dCRY, also known as CRY1-type) serves as a key photoreceptor in brain neurons responsible for clock entrainment (Emery et al. 2000b; Helfrich-Förster et al. 2001; Emery et al. 1998). However, dCRY may also function as a repressor in the peripheral clock of *Drosophila* (Collins et al. 2006). This latter role facilitated its identification in a screen for mutants affecting periodic luciferase oscillations (Stanewsky et al. 1998).

Over time, various methods and approaches have elucidated the molecular mechanisms of the clock, including protein-protein interactions. Classical mutagenesis and genetic mapping were instrumental in identifying the first circadian clock components, while techniques like yeast two-hybrid assays, co-immunoprecipitation, and co-transfection experiments (comparing wild-type and mutant versions) revealed interactions among these components. For example, several key domains have been identified within the dTIM protein: two regions essential for binding PER (PER-binding #1 and PER-binding #2), a cytoplasmic localization domain (CLD), a nuclear localization signal (NLS), and a region where Importin binds (Saez and Young 1996; Jang et al. 2015; Lin et al. 2023) (Figure 1C). The dynamics of PER-dTIM interaction have been studied in great detail using Fluorescence Resonance Energy Transfer (FRET) in *Drosophila* Schneider 2 (S2) cells. While the PER-dTIM interaction is essential for the nuclear translocation of both proteins, once inside the nucleus, either the topology of the proteins or their interaction changes (Meyer et al. 2006; Saez et al. 2011). Additionally, nuclear export signals are critical for the proper functioning of both dTIM and PER (Singh et al. 2019; Giesecke 2023). The stability of dTIM is further regulated by phosphorylation, with shaggy (GSK) and CK2 playing major roles in this process (Top et al. 2016).

**Figure. 1.**
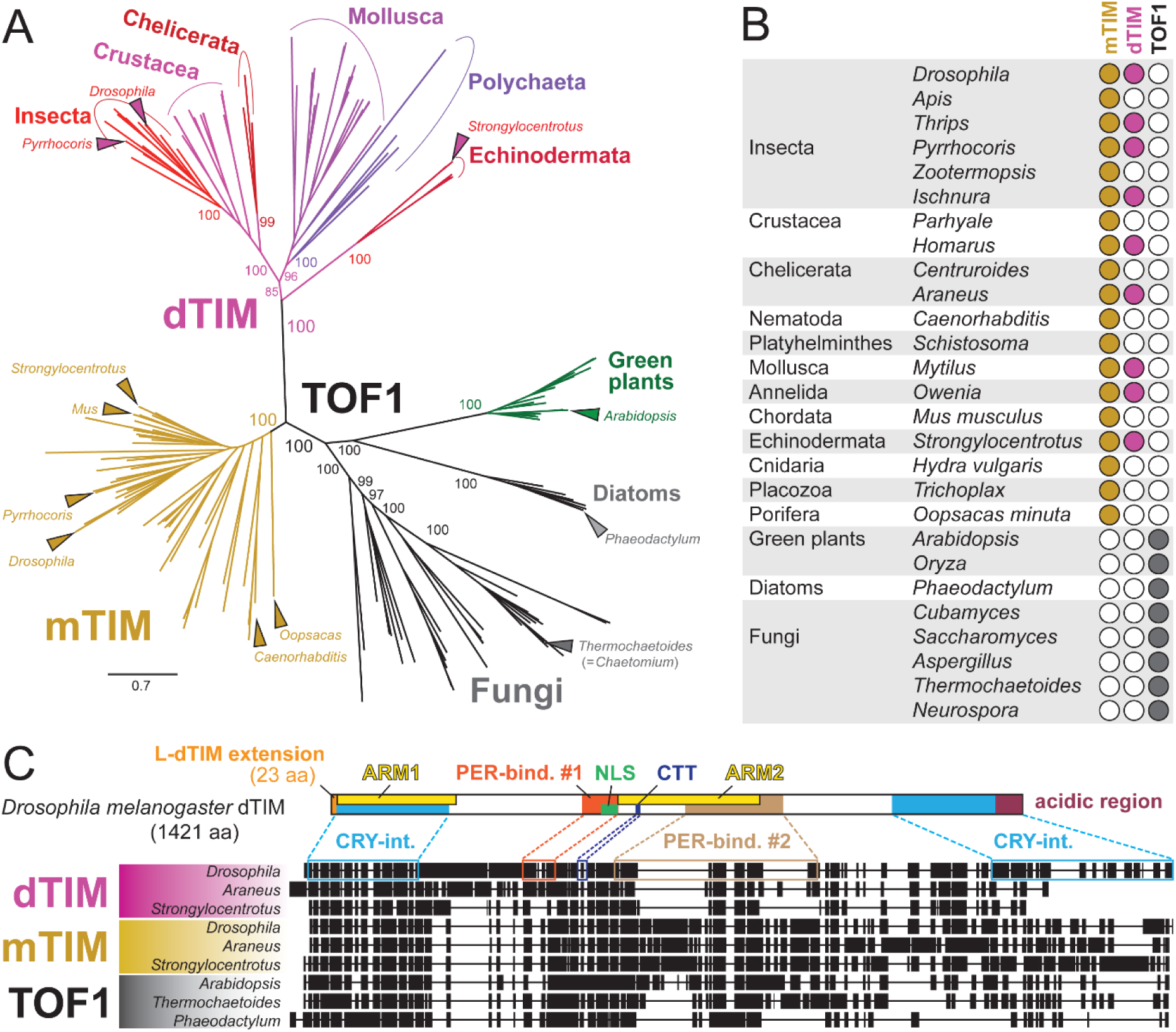
Phylogeny of dTIM (*Drosophila*-type timeless), mTIM (mammalian-type timeless), and Topoisomerase 1-associated Factor TOF1. (A) Phylogenetic tree inferred from protein sequence alignment using RAxML maximum likelihood GAMMA-based model. Major groups are highlighted, with model species indicated by arrowheads. Bootstrap values (inferred from 500 replicates) are shown in %. For the detailed tree including all bootstrap values see Figure S1. (B) The distribution and mutual exclusivity of mTIM and TOF1 suggest that these proteins are orthologs, with mTIM specific to animals and TOF1 to plants, fungi, and diatoms. dTIM is only found in Bilateria, including two basal deuterostomian lineages (Hemichordata, Echinodermata) and most Protostomia. It has been independently lost in chordates, platyhelminths, nematodes, scorpions, termites, hymenopterans, and amphipod Crustacea (*Parhyale*). (*C*) Protein alignment illustrates dTIM-specific deletion located within PER-interaction domain #2 (light brown), conserved from the sea urchin (*Strongylocentrotus*) to *Drosophila*.

The interaction between PER and dTIM is crucial not only for maintaining circadian rhythms under constant conditions but also as part of the entrainment (clock synchronization) mechanism. Since PER is stabilized by dTIM, depletion of dTIM in the morning results in a phase advance, whereas evening depletion triggers a phase delay. Light-mediated degradation of dTIM requires interaction between dTIM, dCRY, and JETLAG (Peschel et al. 2009). Flies with mutated or absent dCRY (Emery et al. 2000a; Dolezelova et al. 2007) or mutated JETLAG (Peschel et al. 2006; Koh et al. 2006) remain rhythmic under constant light—conditions that cause arrhythmicity in wild-type flies. JETLAG, an F-box protein with leucine-rich repeats, targets dTIM for light-induced degradation. Meanwhile, dCRY polyubiquitination and subsequent degradation are triggered by Ramshackle (BRWD3) (Ozturk et al. 2013). Beyond its role in light input, dCRY also contributes to the robustness of rhythmicity under constant conditions at low temperatures in *Drosophila* (Dolezelova et al. 2007). In the monarch butterfly, dCRY depletion reduces eclosion rhythmicity and impairs the robustness of periodic clock gene expression in the brain (Iiams et al. 2024). Similarly, in the silkworm *Bombyx mori*, a dCRY knockout strain exhibited arrhythmic eclosion, and mutant females were unable to properly measure photoperiod (Tobita and Kiuchi 2024).

The role of dTIM in seasonality is supported by studies on a unique mutant in the drosophilid fly *Chymomyza costata*. This species inhabits temperate regions and must synchronize its development with seasonal changes. Long photoperiods promote direct larval development during early instars, whereas short photoperiods induce a developmental arrest known as diapause (Poupardin et al., 2015). Photoperiodic time measurement requires a functional *d-tim* gene, as indicated by mapping a non-photoperiodic diapause mutation to the *timeless* gene. This mutation involves an approximately 3 kb deletion in the promoter region, removing cis-regulatory sequences essential for transcription (Stehlik et al. 2008; Kobelkova et al. 2010). There has been ongoing debate regarding whether the reproductive arrest triggered by low temperature and short photoperiods in *Drosophila melanogaster* constitutes diapause or quiescence. The definition of diapause as an anticipatory mechanism to prepare for harsh conditions (Kostal 2006) suggests the latter. Importantly, *d-tim* alleles significantly influence the incidence of diapause/quiescence (Tauber et al. 2007), and the genetic interaction between *d-tim* and *eya* is involved in regulating this state (Abrieux et al. 2020). Surprisingly, mutations in dTIM’s partner, PER, do not appear to affect reproductive arrest in *D. melanogaster* (Saunders et al. 1989), which contrasts with the synergistic role of these two proteins in the circadian clock. In *Bombyx mori*, dTIM, PER, CLK, and CYC are all critical components of the photoperiodic clock, although the precise mechanism by which they function remains unclear (Tobita and Kiuchi 2022).

For some proteins, such as dCRY, mCRY, CLK, and BMAL1, their structures have been determined using crystallography (Zoltowski et al. 2011; Levy et al. 2013; Michael et al. 2023; Czarna et al. 2013). In the case of PER, only a portion of the protein has been successfully crystallized (Yildiz et al. 2005). However, for other proteins, including dTIM, the 3D structure has remained elusive. While related proteins such as Topoisomerase 1-associated Factor (TOF1) from the fungus *Thermochaetoides* have been partially resolved through crystallography (Grabarczyk 2020), the full structure of dTIM was only recently resolved as part of the dCRY-dTIM dimer using cryogenic electron microscopy (Cryo-EM; Lin et al. 2023). The interaction between dCRY and dTIM involves two distinct regions of dTIM: the amino-terminal Armadillo 1 (ARM1) repeats and a C-terminal CRY-binding domain. Upon light illumination, large-scale rearrangements in dCRY are coupled with conformational changes in its flavin cofactor. This process results in the release of the autoinhibitory C-terminal tail from the dCRY pocket, which is then replaced by the N terminus of dTIM. This terminal motif is highly conserved across dTIM species, and its addition of only 23 amino acids significantly reduces light sensitivity in flies (Rosato et al. 1997; Tauber et al. 2007; Lamaze et al. 2022; Sandrelli et al. 2007; Deppisch et al. 2022).

Interestingly, the paralogous mTIM has been shown to play a role in the circadian clock, albeit to a much lesser extent than dTIM. It influences light sensitivity in *Drosophila*, rhythmicity in crickets and linden bugs, and more recently has been implicated in adjusting activity phases in humans and mice (Benna et al. 2010; Kotwica-Rolinska et al. 2022a; Nose et al. 2017; Kurien et al. 2019). However, the functions of mTIM and TOF1 have generally diverged from those of dTIM. TOF1 directly binds to double-stranded DNA as a component of the large replisome complex (Baretic et al. 2020), while mTIM stabilizes replication forks, pairs sister chromatids, and regulates the S phase of the cell cycle (Unsal-Kacmaz et al. 2005). Moreover, despite the overall structural similarities among dTIM, mTIM, and TOF1, specific details appear to be unique to each protein type (Lin et al. 2023).

Despite the relatively well-characterized role of dTIM in the *Drosophila* clock, its function in other organisms appears to be only partially conserved. In vertebrates and several arthropod lineages, dTIM and dCRY have been completely lost (Rubin et al. 2006; Kotwica-Rolinska et al. 2022a), with mCRY serving as the partner of PER in these species (Yuan et al. 2007). Furthermore, vertebrates have undergone multiple rounds of genome duplication, resulting in the multiplication of many clock components, including PERs and mCRYs. In non-*Drosophila* insects, various combinations of mCRY, dTIM, JET, and FBXL3 have been identified (Kotwica-Rolinska et al. 2022a; Deppisch et al. 2023; Dolezel 2023). Functional experiments suggest that the roles of these components in the pacemaker system of some species differ significantly from those in *Drosophila*. The non-essentiality of certain components for clock rhythmicity, as demonstrated for dTIM in the linden bug *Pyrrhocoris apterus*, cricket, firebrat, and cockroaches, suggests potential transitions from one clock setup to another (Kotwica-Rolinska et al. 2022a; Kamae et al. 2012; Danbara et al. 2010; Werckenthin et al. 2020).

The growing availability of transcriptomic and genomic data from all major animal lineages has enabled systematic analyses of the evolution of specific clock proteins. In this study, we focused on a detailed comparison of dTIM, aiming to identify its origin and map its occurrences and losses across the animal phylogeny. We analyzed conserved and variable regions of the protein, evaluating them in the context of *d-tim* gene evolution, including its exon/intron structure. Additionally, we examined the coevolution of dTIM with its partner proteins—PER, dCRY, mCRY, and JETLAG—and, by mapping these changes onto the animal phylogeny, highlighted potential key interactions that have shaped the evolution of the animal circadian clock. Finally, we investigated alternative splicing of *d-tim* in Diptera, proposing conserved molecular mechanisms that may regulate dTIM in this insect order.

## RESULTS

### dTIM Is an ortholog of animal mTIM and TOF1 from plants, fungi, and diatoms

We systematically searched for dTIM, its related mammalian-type TIM (mTIM), and TOF1 proteins across diverse organisms and reconstructed the phylogeny of the identified sequences (Figure 1A; Figure S1). Consistent with previous reports, dTIM was identified in most Protostomia and two basal deuterostome groups, Echinodermata and Hemichordata. For the latter group, however, dTIM was only found in the acorn worm *Ptychodera flava*, while it was absent in another hemichordate, *Saccoglossus kowalevskii*. Notably, we did not retrieve dTIM from any Cnidaria, Placozoa, or Porifera, suggesting that dTIM likely originated at the dawn of Bilateria.

The paralogous mTIM, a protein essential for development and circadian clock function in mammals and insects (Gotter et al., 2000; Benna et al., 2010; Barnes et al., 2003), is found in all animals, including “prebilaterian” groups such as Cnidaria, Placozoa, and Porifera. Furthermore, the occurrence of mTIM is mutually exclusive with that of TOF1, a protein found in plants, diatoms, and fungi (where it is known as Swi1 in *Schizosaccharomyces pombe*). This suggests that mTIM and TOF1 are one-to-one orthologs (Figure 1B). Protein alignment reveals conserved dTIM-specific deletions in the region containing PER-binding domain #2 (PER-bind. #2), which clearly distinguishes dTIM from mTIM (Figure 1C). The similarity among mTIMs and TOF1 proteins is highest within the first ∼600 amino acids, whereas the C-terminal region is notably variable.

### Independent losses of dTIM

In addition to the well-supported and previously described losses of dTIM in Chordata, Hymenoptera, and termites (with the possible exception of early-branching termite species such as *Porotermes*) (Rubin et al., 2006; Kotwica-Rolinska et al., 2022a), we did not identify this gene in Amphipoda (Crustacea), Scorpiones (Chelicerata), Platyhelminthes, or Nematoda (Figures 1B and 2F; see Figure S1 and Table S1 for details). The *Caenorhabditis elegans* TIM homolog TIM-1, previously analyzed in the context of chronobiology (Hasegawa et al., 2005), belongs to the mTIM clade (Figure 1A).

**Figure 2.**
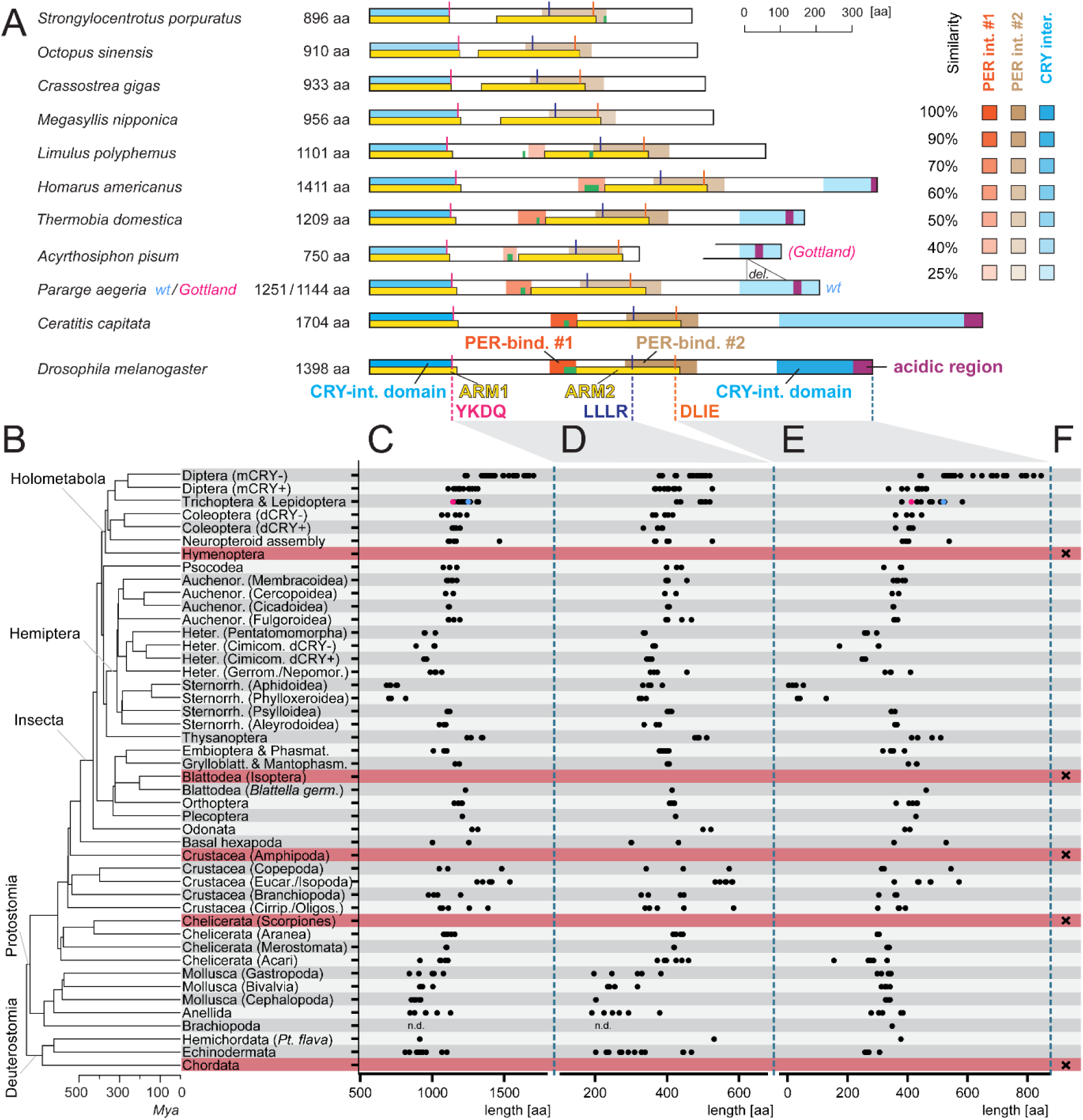
Differences in dTIM proteins. (A) Selected examples of dTIM proteins with functional domains annotated based on sequence similarity, highlighting major variations. Shades indicate similarities in PER-binding and CRY-interaction domains. Two alleles in *Pararge aegeria* differ in the CRY-interaction domain. For additional protein models and similarity details, see Figure S2 and S5. (B) Phylogenetic relationships of bilaterian groups analyzed for dTIM. Lineages where dTIM has likely been lost are shaded in brick-red. (C) Total dTIM length, (D) the central segment (YKDQ to LLLR), and (E) the terminal portion (DLIE motif to C-terminus) are plotted, with each dot representing one protein from a species in a particular taxonomic group. *P. aegeria* alleles are marked in magenta and turquoise, corresponding to panel A. (F) dTIM loss is indicated by “x.” The *Lingula* (Brachiopoda) sequence is incomplete at the N-terminus; thus, measurements were not determined (*n.d.*).

### Variability in dTIM

The available dataset allowed us to explore the sequence variability of dTIM proteins. The length of dTIM ranged from 683–757 amino acids (aa) in aphids and 874–920 aa in cephalopod mollusks, reaching up to 1666–1704 aa in the Tephritidae family of Diptera (including *Ceratitis* and *Bactrocera* genera) (Figure 2C, Table S2). Since *Drosophila* dTIM is by far the best-characterized member of this group—with extensive functional assays (Saez and Young, 1996; Jang et al., 2015), genetic mutant studies (Rothenfluh et al., 2000a, 2000b; Matsumoto et al., 1999; Singh et al., 2019), and structural analyses (Lin et al., 2023)—we used it as a reference for domain and region terminology in our interspecies comparisons.

The most conserved regions across species are the two armadillo domains (Figure 2A). The first domain, ARM1, includes a region that interacts with the importin 1 protein, where several mutations impacting the free-running period have been reported (Jang et al., 2015; Top et al., 2016; Thakkar et al., 2023). ARM1 is nearly identical to the first CRY-interaction domain, whereas the second armadillo domain (ARM2) partially overlaps with PER-binding site #2 (PER-bind. #2), which was functionally identified using *Drosophila* S2 cell cultures (Saez and Young, 1996).

In contrast, PER-binding site #1 (PER-bind. #1), located between the two armadillo domains, exhibits greater sequence variability and can be detected in insects, Crustacea, and, to some extent, chelicerates. The absence of PER-bind. #1 correlates with a shorter central part of the protein in certain Mollusca and basal Deuterostomia (Echinodermata, Hemichordata). In these groups, only the acidic region typical of the N-terminal part of PER-bind. #1 is partially conserved (Figure 2, Figures S2, S3, and S4). Additional variability in the central region (between the armadillo domains) is particularly pronounced in some Crustacea, an ancient and highly diverse group from which Hexapoda (including insects) evolved.

The most variable part of dTIM proteins is located in the C-terminal region. In *D. melanogaster*, this region contains a CRY-interaction domain and a C-terminal acidic region responsible for cytoplasmic localization, also known as the cytoplasmic localization domain (CLD) (Saez and Young, 1996; Lin et al., 2023; Cai et al., 2021). This region has notably expanded in a subset of Diptera, including *Drosophila* and *Ceratitis*, whereas the shortest sequences are found in aphids and phylloxera, which differ dramatically from their sister group, Psylloidea (Figure 2E). Remarkably, the geographical allele Gottland of the Speckled Wood (*Pararge aegeria*) is characterized by a substantial deletion (Lindestad et al., 2022) that encompasses the CRY-interaction domain (Figure 2A, E), suggesting that this allelic variant may result in altered light sensitivity. The CRY-interaction domain is recognizable even in the most ancestral insect, *Thermobia*, and is detected in some, but not all, Crustacea (Figures S2 and S4). Interestingly, variability among paralogs within individual species—most prominently in the *Daphnia* genus, which has up to 8–11 paralogs—centers on the inter-armadillo region and the C-terminal tail. These regions are also the most variable in interspecific comparisons (Figures 3 and S5).

**Figure 3.**
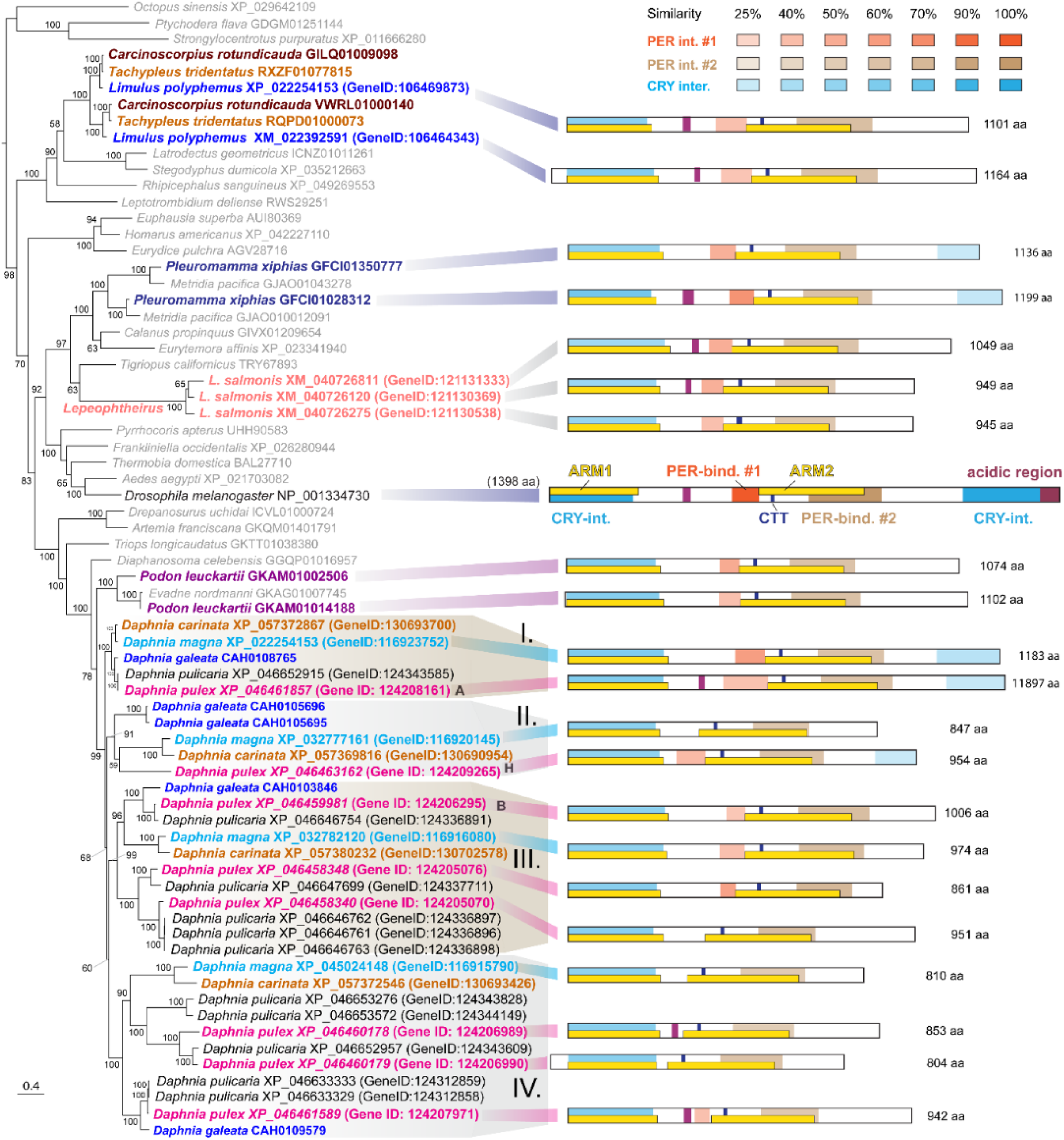
Independent dTIM gene dupliplications were detected in horseshoe crabs (*Limulus*, *Tachypleus*, and *Carcinoscorpinus*), copepods (*Pleuromamma*, *Metridia*, and *Lepeophtheirus*), and branchiopods (*Podon* and *Daphnia*). The phylogenetic tree inferred using RAxML and Bootstrap values (500 replicates, shown in %), indicates these duplications are lineage specific. In the *Daphnia* genus, four primary dTIM clusters (I.-IV.) are present across all species. Additional species-specific duplications resulted in 8 paralogs in *D. pulex* and 11 in *D. pullicaria*. Protein models show conserved similarities in PER-binding and CRY-interaction domains. Labels A, B, and H for *D. pulex* follow the terminology of Bernatovicz et al. (2016).

### Independent duplications of dTIM

In several crustacean and chelicerate species, multiple *d-tim* paralogs were identified. While two paralogs were most commonly observed, *Daphnia* species exhibited varying numbers of paralogs, reaching as many as 11 in *D. pullicaria*. To clarify the origin of these duplications and assess whether *d-tim* gene duplication could contribute to a more general pattern in dTIM evolution, we performed a detailed phylogenetic analysis. As shown in Figure 3 and well supported by bootstrap analysis, *d-tim* duplication can be mapped to only five lineages. In horseshoe crabs (*Limulus polyphemus*, *Carcinoscorpius rotundicauda*, and *Tachypleus* tridentatus; Figure S6), two *d-tim* paralogs are consistent with the genome duplication events in this lineage, which includes also circadian transcription factor BMAL1 and other bHLH-PAS proteins (Tumova et al. 2024). In copepods (Crustacea), three closely related *Lepeophtheirus salmonis* paralogs (Figure S7) resulted from duplications independent on two paralogs found in *Pleuromamma* and *Metridia*. In branchiopods, the duplication that produced two *d-tim* genes in *Podon* occurred independently of the series of duplications specific to *Daphnia* species. Four primary dTIM clusters (I–IV) are present across all species in the *Daphnia* genus (Figure 3). The longest proteins are found in cluster I, which branches at the base of the *Daphnia*-specific tree expansion. Proteins from cluster I also contain a complete set of functional domains. Furthermore, the *Daphnia pulex* dTIM paralog A, belonging to cluster I, is cyclically expressed (Bernatovicz et al., 2016). Clusters II, III, and IV include additional species-specific duplications, resulting in eight paralogs in *D. pulex* and eleven in *D. pullicaria*.

### Substitution rate in dTIM and coevolution with partner proteins

To provide a clear graphical representation of protein sequence variability, we plotted the substitution rate per amino acid position per million years for each analyzed dTIM protein. This method is minimally influenced by in-frame deletions or variations at phylogenetically variable positions, as evidenced by the minimal differences in substitution rates between mCRY+ and mCRY- Diptera (Figure 4B), despite these groups showing significant variation in the length of their C-terminus (Figure 2E). However, this approach effectively highlights taxa with alterations in phylogenetically conserved positions, such as aphids and phylloxera. Notably, the lowest substitution rates are observed in cephalopods, a group of mollusks that includes octopuses, cuttlefish (*Sepia*), and squid.

**Figure 4.**
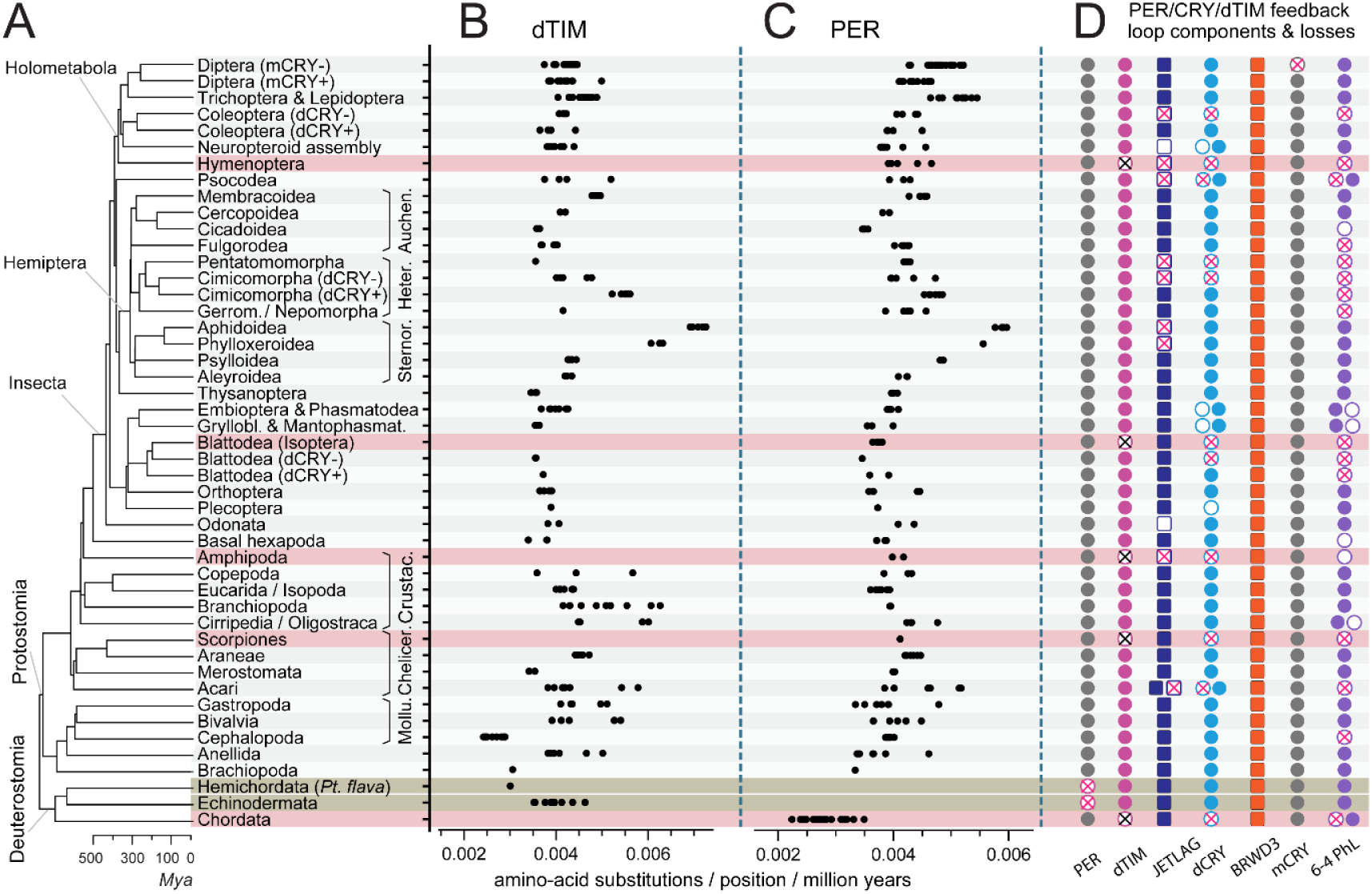
Coevolution of dTIM with partner proteins. (A) Phylogenetic relationships among the analyzed bilaterian groups. Protein changes quantified as substitutions per position per million years, shown for (B) dTIM and (C) PER. Each dot represents a protein from a species within a specific higher taxon. (D) Presence (filled circle), absence (empty circle), and loss (x) of key components in the PER/CRY/dTIM negative feedback loop across lineages. Loss of dTIM is highlighted by a light red background, while PER loss (in Hemichordata and Echinodermata) is marked with a grey-brown background.

To further explore the potential causes of dTIM variability, we examined dTIM-interacting proteins. PER, a key partner in the *Drosophila* clock, is present in nearly all Bilateria, with the unique exceptions of two basal deuterostomian groups: Hemichordata and Echinodermata. Notably, even the loss of PER did not affect the substitution rate of dTIM. Conversely, the highest substitution rates observed in dTIM from aphids and phylloxera were mirrored by increased variability in PER within these species, suggesting coevolution of the two proteins (Figure 4C, D).

The loss of dTIM is consistently accompanied by the loss of dCRY, as observed in Hymenoptera, Isoptera, Amphipoda, Scorpiones, and Chordata (Figure 4D). However, several instances exist where dCRY has been independently lost while dTIM remains present. These cases include certain Coleoptera (most Polyphaga, including the red flour beetle *Tribolium*), a subset of Blattodea (including *Periplaneta americana*, Bazalova et al., 2016), and a subset of Heteroptera. Notably, within Heteroptera, dCRY has been lost at least twice independently, as evidenced by its complete absence in the order Pentatomomorpha and parts of Cimicomorpha (family Cimicidae). This conclusion is further supported by synteny analysis. For example, *Lethocerus indicus* (Nepomorpha), representing an ancestral lineage of Heteroptera, retains the *d-cry* gene. Comparative analysis of the *d-cry* locus in *Lethocerus* and *Riptortus pedestris* suggests that *d-cry* loss in the latter species resulted from a combination of several chromosomal inversions. In contrast, different chromosomal rearrangements led to the loss of *d-cry* in the genus *Cimex*. Meanwhile, the sister lineage of Cimicomorpha, Miridae, still retains the *d-cry* gene (Figure 5A, B, and Figure S8A).

**Figure 5.**
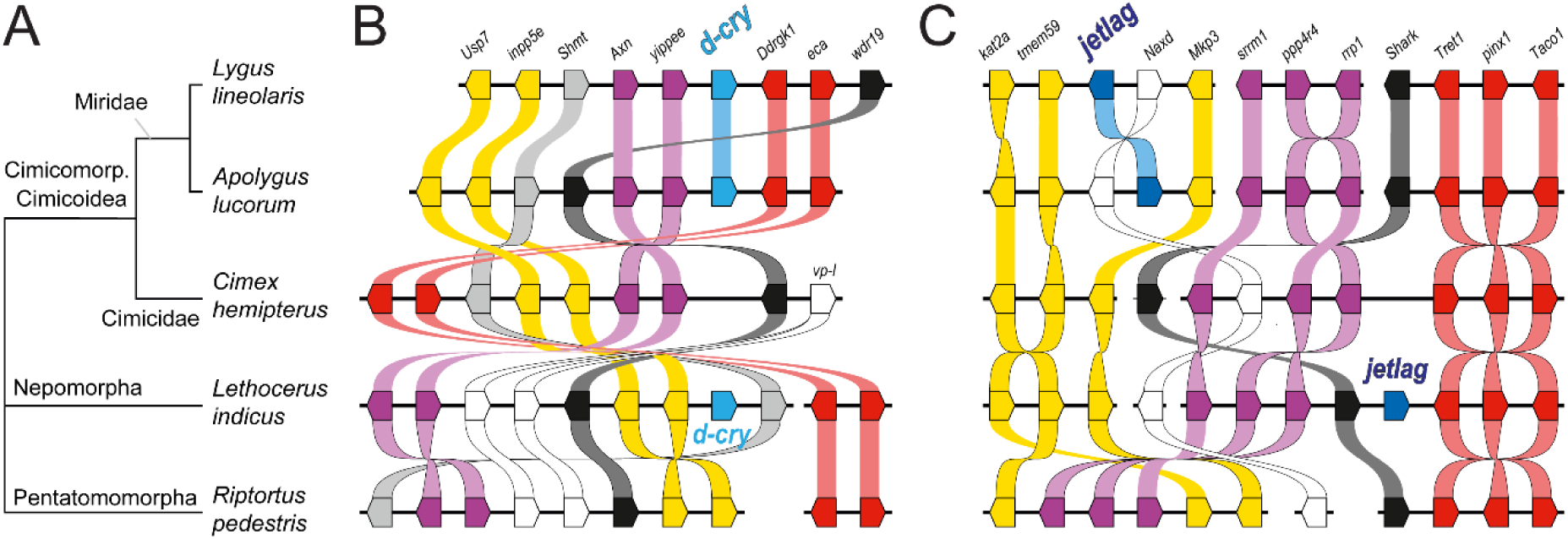
Gene synteny illustrates the independent loss of *Drosophila*-type *cryptochrome* (*d-cry*) and *jetlag* (*jet*) genes in *Cimex hemipterus* and *Riptortus pedestris*. (A) Phylogenetic relationship among analyzed five (pan)heteropteran species is presented alongside (B) *d-cry* and (C) *jet* gene syntenies. The genomic positions of *d-cry* and *jet* in *L. lineolaris*/*A. lucorum* (Miridae) and *L. indicus* (Nepomorpha) differ, as indicated by distinct sets of neighboring protein-coding genes. Although synteny genes can be mapped to *C. hemipterus* (Cimicidae) and *R. pedestris* (Pentatomorpha), their composition and orientation vary, supporting independent losses of *d-cry* and *jet*. Horizontal black lines represent genomic contigs/scaffolds (not to scale). Arrow-like boxes indicate protein-coding genes and their orientation. Orthologous genes are color-coded and interconnected. For detailed synteny depictions, see Figure S8 and Tables S3 and S4.

Our analysis reveals that the loss of dCRY is consistently accompanied by the loss of JETLAG in several taxa, including Hymenoptera, a subset of Coleoptera, Phthiraptera (a subset of Psocodea), Amphipoda, Parasitiformes (a group within Acari that includes ticks), and a subset of Heteroptera. Remarkably, the two independent losses of dCRY in Heteroptera (Pentatomomorpha and Miridae) are also associated with the loss of the *jetlag* gene. Gene synteny analysis suggests that independent chromosomal rearrangements underlie these losses of *jetlag* in Pentatomomorpha and Cimicidae (Figure 5C and Figure S8B). Despite the loss of dCRY, the organization of dTIM domains (Figure S2) appears to be very similar among *Lygus* (where dCRY is still present), *Cimex* (where dCRY has been lost), and *Pyrrhocoris* (where dCRY has been independently lost). Notably, dTIM in the early-branching species *Lethocerus indicus* differs significantly from these species as it retains an elongated CRY- interaction domain at its C-terminus.

The losses of *d-cry* and *jetlag* do not align perfectly across all bilaterian species. For instance, *jetlag* (annotated as *F-box and LRR protein 15* in chordates) is retained in Chordata and Blattodea (Figure 4D). Termites (Isoptera), a subset of Blattodea that have progressively lost *dCRY* and even *dTIM*, still possess *jetlag*. Interestingly, a unique and substantial modification of *dTIM* in Aphidomorpha (Phyloxeroidea and Aphidoidea) is not associated with the loss of *d-cry*. The *d-cry* gene is present in all Aphidomorpha species, and dCRY protein was detected in the brains of the pea aphid *Acyrthosiphon pisum* via immunohistochemistry (Colizzi et al., 2021). This suggests that the rapid evolution of dTIM in Aphidomorpha is lineage-specific, as their sister groups, Psylloidea and Aleyroidea, possess dTIM with a generally conserved domain structure (Figure S2). Notably, the loss of *jetlag* corresponds precisely to the accelerated accumulation of changes in *dTIM* within Aphidomorpha. Moreover, the PER protein also shows an elevated substitution rate in this lineage (Figure 4).

### Changes in gene structure underlie dTIM evolution

To gain further insight into *dTIM* changes, we compared the *d-tim* gene structure across selected representative species. Even visual inspection revealed a complex remodeling of *d-tim* exons in most Crustacea, with only the early-branching *Darwinula* showing a structure similar to both the horseshoe crab *Limulus* (Chelicerata) and the firebrat *Lepisma* (Insecta). Similarly, exon fusion in *Orchesella* (Collembola) appears to be lineage-specific, which is why these species are presented in a separate box (Figure 6). Importantly, a generally conserved exon pattern is observed in the majority of the species analyzed, with several exons traceable from echinoderms to insects. Consistent with the variability in dTIM protein, the most conserved regions encompass the exons encoding the ARM1 and ARM2 domains. In the case of ARM1, a Holometabola-specific fusion of four exons is observed, while a similar, though less pronounced, exon fusion can be identified in ARM2.

**Figure 6.**
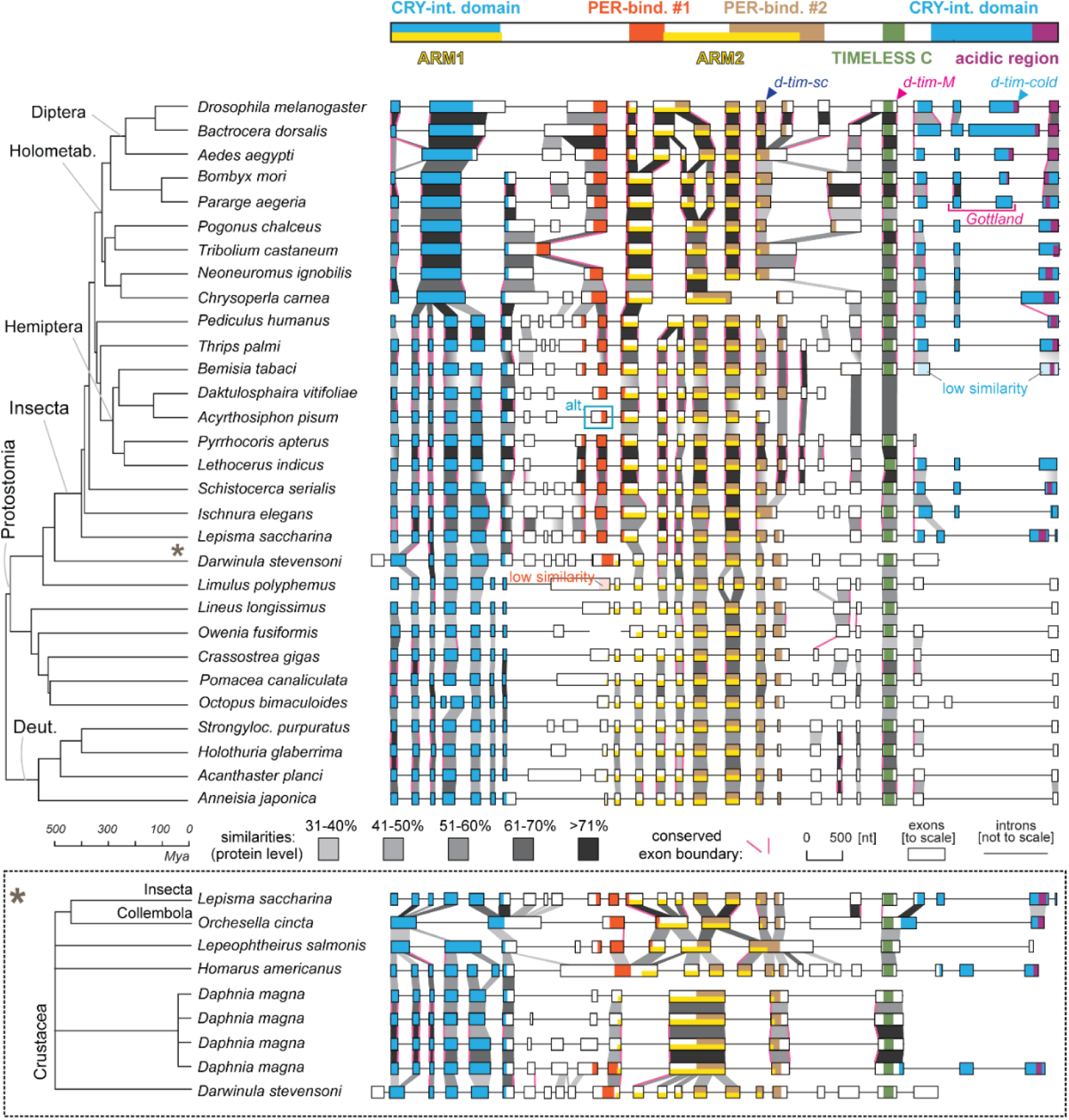
Gene structures and protein domains of dTIMs in Bilateria. Introns, exons, and homologous regions illustrate the conserved and variable parts of *d-tim* gene. Vertical blocks in shades of grey represent sequence similarities at the protein level, while vertical magenta lines indicate conserved exon boundaries. Protein domains and functional regions are annotated based on their similarity to *Drosophila* dTIM (shown at the top). For clarity, the complexity specific to Crustacea is presented in a separate box (*), where *Lepisma* and *Darwinula* appear in both analyses. The highest variability is observed in the PER-binding region #1 and the CRY-interaction domain, located at the C-terminus. Within Insecta, the CRY-interaction domain has been lost in *Acyrthosiphon pisum* and *Daktulosphaira vitifoliae* and independently in *Pyrrhocoris apterus*. In *Pararge aegeria*, a unique geographical allele is the result of a deletion spanning two exons (highlighted with a magenta horizontal line). Thresholds for plotting specific regions were as follows: similarities greater than 45% for ARM1, ARM2, and PER- binding regions (except for *Limulus*, where similarity is 38%), and greater than 25% for the CRY- interaction domain (with the lowest similarity observed in *Bemisia*, at 19%). Intron retention in three *Drosophila* splicing isoforms is marked with arrowheads, and alternative exon removal in *A. pisum* (Barbera et al., 2017) is indicated by a turquoise rectangle with a note “alt”.

The dTIM region between the ARM1 and ARM2 domains exhibits significant variability. Notably, some level of similarity is detected in insects, particularly in the PER-binding #1 region; however, beyond insects, this portion of dTIM shows high divergence. Similarly, the exons encoding this region are only minimally conserved within insects and display no detectable similarity in Deuterostomia or ancestral Protostomia, including lineages such as Mollusca, Nemertea, Polychaeta, and Chelicerata. Within insects, the variability between the ARM1 and ARM2 domains is most pronounced in Aphidomorpha, where similarity falls below the plotted 30% threshold. This highlights the remarkably high degree of modification of dTIM in both Phylloxeroidea and Aphidoidea (Figure 6 and Figure S3). Furthermore, early work identified alternative splicing of the *d-tim* exon encoding the first part of the PER-binding #1 region in two aphid species (Barbera et al., 2017); this splicing is distinct from several alternative splicing isoforms detected in *D. melanogaster* (Figure 6, top).

The C-tail of dTIM, located downstream of the ARM2 domain, contains a few conserved exons. In *D. melanogaster*, a Drosophilidae-specific fusion to the upstream exon has been detected (Figure 7). This region encodes the TIMELESS C domain (Pfam: PF05029, Figure S9), where similarity is observed even between dTIM and mTIM (Koike et al., 1998). When mutated, various degrees of temperature-dependent changes in the free-running period have been described (Matsumoto et al., 1999; Singh et al., 2019). Interestingly, a major truncation in Aphidomorpha removes the C-tail and even the otherwise highly conserved TIMELESS C domain (Figure 6).

**Fig. 7.**
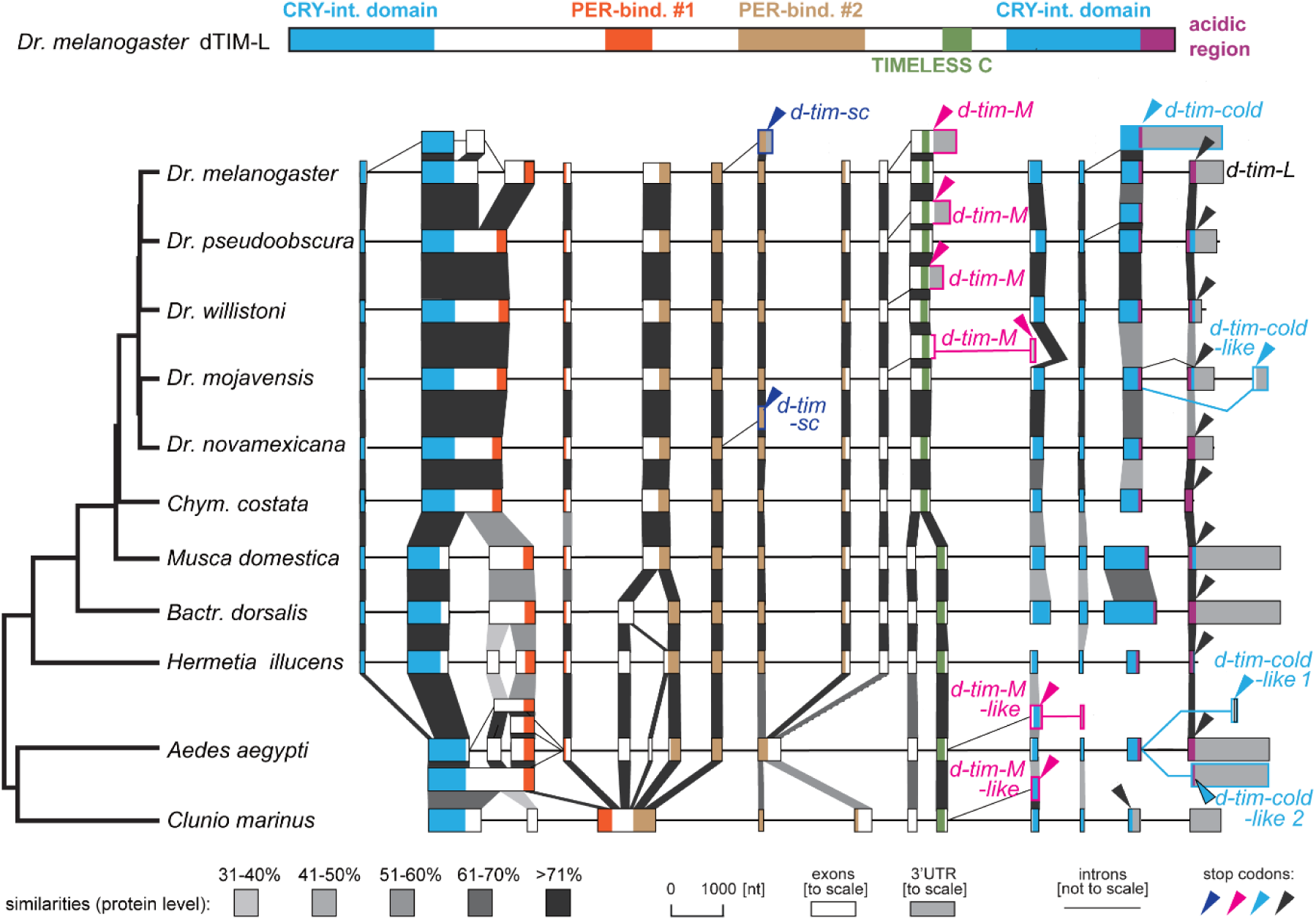
*d-tim* gene structures in Diptera with highlighted alternative splicing isoforms. The similarities and dTIM domains are depicted identically as in Figure 6. Four major splicing isoforms exist in *D. melanogaster*: The longest isoform is the canonical *d-tim-L*. Another isoform, *d-tim-cold* (turquoise), is produced by retention of the last intron at low temperatures, resulting in the absence of the last exon. At high temperatures, intron retention generates the *d-tim-M* (magenta) isoform. Additionally, the *d-tim-sc* (blue) isoform is abundant at low temperatures. Identical or comparable isoforms have been observed in some other dipteran species. For example, modified exon splicing produces *tim- cold*-like isoforms in *D. mojavensis* and *Aedes*. In the case of *d-tim-M*-like isoforms in *Aedes* and *Clunio*, the intron downstream of the *D. melanogaster* M isoform is spliced. Despite this difference, the effect on *d-tim* mRNA destabilization may be similar to the impact of *d-tim-M* in *D. melanogaster*.

The high variability in the protein tails downstream of the TIMELESS C domain is evident at the gene structure level, where only two exons are detected in Deuterostomia (*Strongylocentrotus* and *Acanthaster*), Mollusca (*Crassostrea*), Annelida (*Owenia*), and Chelicerata (*Limulus*). In insects, expansion of the tail correlates with similarities to the CRY-interaction domain in *D. melanogaster*. This domain has been independently lost in *Pyrrhocoris apterus* and Aphidomorpha. The latter is a sister group to *Bemisia tabaci*, a species in which only low similarity (19%) to the CRY-interaction domain is detected (Figures 6 and S3). A complete C-tail is found in *Lethocerus indicus*, a sister taxon to *P. apterus*, where truncation removes the entire CRY-interaction domain downstream of the TIMELESS C domain. In Lepidoptera, gene analysis of the Speckled wood butterfly (*Pararge aegeria*) reveals that the loss of two entire exons in the Gottland allele results in a substantial in-frame deletion in the CRY-interaction domain. In contrast, the expansion of dTIM in Diptera lacking mCRY (mCRY-Diptera) is due to exon extensions, as illustrated in *Bactrocera dorsalis* and *Musca domestica* (Figures 6 and 7).

### N-terminal extensions are rare in dTIM proteins

Gene model analysis suggests that *Darwinula* d-tim may encode a protein extended by 61 amino acids at the N-terminus. Similar N-terminal extensions were found in *Macrobrachium* (Crustacea) and some dTIM paralogs of *Daphnia pulex* and *Limulus* (Supplementary Figures S3 and S5). Importantly, even a shorter 23 amino acid extension in *D. melanogaster* has a significant impact on its interaction with dCRY, resulting in lower sensitivity to light, including altered behavioral rhythmicity under constant dim light (Lin et al. 2023; Sandrelli et al. 2007; Deppisch et al. 2022; Lamaze et al. 2022). In *D. melanogaster*, the N-terminal extension is encoded by an in-frame alternative start codon in the *ls-tim* allele (Rosato et al. 1997). Since this allele is believed to be of recent origin, specific to *D. melanogaster* (Tauber et al. 2007), it was surprising that BLAST-P identified several L-dTIM protein sequences in GenBank for various *Drosophila* species (*D. ananassae* EDV31195; *D. kikkawai* XP_041631545). However, closer inspection revealed that these protein sequences are clear annotation errors, as the available genomes only encode short (s-dTIM) proteins.

### Conserved features of the alternative splicing pattern in Diptera

Alternative splicing is a key regulatory mechanism of dTIM in *D. melanogaster* at both low and high temperatures (Boothroyd et al. 2007; Shakhmantsir et al. 2018; Martin Anduaga et al. 2019; Foley et al. 2019; Montelli et al. 2015). The combination of genomic sequences and available transcriptome shotgun assemblies (TSAs) allowed us to investigate whether a similar splicing pattern exists in other Diptera (Figure 7). The *d-tim-sc* (*tim-short and cold*), resulting from an alternative polyadenylation site, was identified only in *D. novamexicana* and confirmed in *D. melanogaster*.

A second low-temperature-dependent regulation of *d-tim* in *D. melanogaster* is the retention of the last intron (located between the penultimate and final exons), which results in a premature stop codon that removes the acidic region at the C-terminus. This portion of the dTIM protein is phosphorylated by Casein Kinase 2, which in turn inhibits nuclear export and impacts CLOCK transcriptional activity (Cai et al. 2021). Although the last intron is generally conserved across several species, splicing was minimally affected in *Musca* (Bazalova and Dolezel 2017). However, we identified two distinct alternative splicing patterns in the mosquito *Aedes aegypti* and the fly *D. mojavensis* that either remove or modify the C-terminal part of dTIM. In the first pattern, an alternative splice site within the exon removes a portion of the protein. In the second, the penultimate canonical exon is fused with an alternative last exon located at the very end of the gene, resulting in an altered amino acid sequence. Since both splicing patterns were observed in *Aedes aegypti* and *D. mojavensis*, and one of them was confirmed in *Clunio marinus*, this suggests that the mechanism may be conserved.

The last major dTIM isoform, *medium* (*d-tim-M*), is expressed at high temperatures and is characterized by the retention of an intron in the center of the mRNA. The resulting transcript does not appear to be translated; instead, this mechanism reduces the pool of dTIM-coding mRNAs via nonsense-mediated decay (NMD) (Martin Anduaga et al. 2019). A comparable splicing pattern was identified in three *Drosophila* species, with a slight modification observed in *D. mojavensis*. Interestingly, in *Aedes* and *Clunio*, alternative splicing is detected one exon downstream of the canonical *d-tim-M* isoform (Figure 7). Although no experimental data are available, and it is unknown how temperature affects this splicing pattern, the similarity with the *d-tim-M* isoform suggests that a potentially comparable regulatory mechanism might also be present in *Clunio* and *Aedes*.

## DISCUSSION

Although the major role of circadian clocks—measuring 24 hours—is conserved among organisms, the molecular machinery achieving this task appears to be variable among Bilateria, a group relying on homologous clock genes. This study focused on the variability and evolution of one of these proteins, dTIM, by exploiting available genomic and transcriptomic data (see Figure 8 for a summary). While dTIM originated from mTIM in early Bilateria, its functional participation in the circadian clock is confirmed in only several insect species representing the orders Diptera, Heteroptera, Orthoptera, and Zygentoma. However, its role varies even among these groups. In *Drosophila*, dTIM is an essential clock component, whereas in other species, such as the linden bug *P. apterus*, the cricket *Gryllus bimaculatus*, and the firebrat *Thermobia domestica* (which belongs to the most ancestral insect group, Zygentoma), it serves as a mere modulator of clock pace. A key feature of dTIM in *Drosophila* is its interaction with the PER protein, which was mapped in the seminal study using the *Drosophila Schneider 2* cell line (Saez and Young 1996). Out of the two identified PER-binding sites, one is conserved in dTIMs across all studied lineages. Since a substantial part of this site overlaps with the ARM2 domain, it is difficult to determine which of these functions is responsible for the high conservation. The other site, PER-bind #1, is well conserved in Insecta and some Crustacea (although the latter also contain species with variable and divergent dTIM sequences), and it is still identified in Chelicerata, albeit with lower similarity.

**Figure 8.**
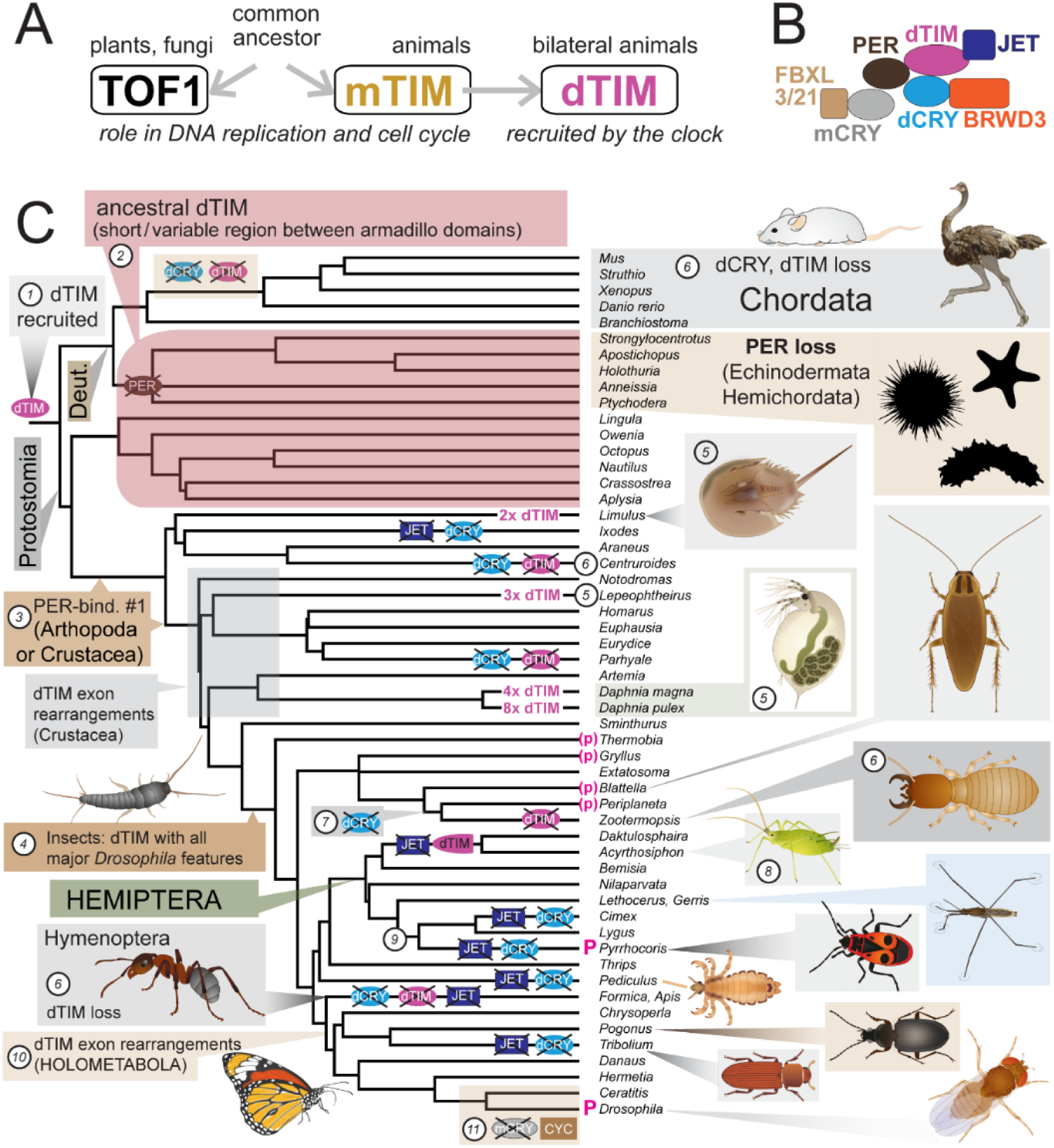
Summary of major changes in dTIM evolution. (*A*) dTIM originated from mTIM by gene duplication in the common ancestor of Bilateria. (*B*) Schematic depiction of key components in the negative PER/mCRY/dTIM feedback loop in circadian clocks. (*C*) Evolutionary changes in dTIM and its interacting clock components were mapped onto the phylogeny of Bilateria: (1) dTIM was recruited in the common ancestor of Protostomia and Deuterostomia and (2) dTIM is retained in ancestral lineages of both Protostomia and Deuterostomia. (3) The PER-binding domain #1 evolved either in Arthropoda or Crustacea. (4) All *Drosophila*-specific features first appeared in ancestral insect dTIM and these key features are retained in dTIM of most insects. (5) Lineage-specific duplications resulted in multiple dTIM paralogs in *Limulus* (and other horseshoe crabs), *Lepeophtheirus*, and *Daphnia*. (6) dTIM has been lost in several lineages, including Chordata, scorpions (*Centruroides*), Amphipoda (*Parhyale*), Hymenoptera, and Isoptera (*Zootermopsis*). These losses consistently coincide with the absence of dCRY. (7) In *Blattodea*, the loss of dCRY precedes the loss of dTIM in termites. (8) Changes in the dTIM sequence in aphids and *Phylloxera* correlate with the loss of JET. Independent loss of dCRY and JET occurred in two Heteroptera lineages (9), suggesting a mutation in their common ancestor that impacted dCRY and JET function in the circadian clock. (10) A major fusion of *d-tim* exons is observed in Holometabolous insects. (11) The loss of mCRY and simultaneous modification of BMAL1 into CYC dates to Cyclorrhapha (e.g. *Drosophila*). Species with circadian clock phenotypes assessed from *d-tim* genetic mutants are marked with a magenta “P.” Phenotypes derived from *d-tim* RNAi silencing experiments are indicated with “(p).”

In Annelida and Mollusca, the protein region corresponding to PER-bind #1 is substantially modified, inviting us to speculate whether dTIM even interacts with PER in these organisms. In Hemichordata and Echinodermata, two lineages that lost PER completely, the level of modifications in the region between ARM1 and ARM2 is comparable to that detected in Mollusca and Annelida. Furthermore, low substitution rates per amino acid position are observed in dTIM of Mollusca, Annelida, Hemichordata, and Echinodermata. Thus, even the absence of PER had minimal impact on dTIM evolution in early-branching Deuterostomia.

A second role of dTIM, light-mediated entrainment, is documented in great detail in *D. melanogaster*, where the light-triggered dTIM-dCRY interaction—elegantly described in the yeast system, later in *Drosophila* S2 cells and flies, and most recently by Cryo-EM (Ceriani et al. 1999; Busza et al. 2004; Lin et al. 2023)—results in ubiquitination and subsequent degradation of both proteins, a process in which JET and BRWD3 play a key role (Peschel et al. 2009; Ozturk et al. 2013). Except for BRWD3, a gene essential during development (D’Costa et al. 2006), multiple well-supported cases of gene loss are reported for dTIM, dCRY, and JET. Usually, but not always, several components are lost simultaneously. For example, the absence of dTIM is always accompanied by the loss of dCRY (5 independent cases).

However, the absence of dCRY does not always coincide with the loss of dTIM. A simultaneous absence of dCRY and JET is reported in 7 independent cases. These examples of gene losses suggest that once one of the components is missing, the system is compromised, and additional components are often quickly lost. Particularly intriguing is the independent loss of both genes in *Pentatomomorpha* and *Cimicidae*. Notably, the absence of dCRY and JET aligns with the relatively small impact of light as the entraining cue in the linden bug *P. apterus* (Kaniewska et al. 2020). It will be interesting to determine the light entrainment capacity in Miridae, a lineage where dCRY and JET are still present. In general, RNA interference seems to work reliably in true bugs, making it possible to explore the specific roles of these proteins. The loss of dCRY and JET is typical for a large portion of Coleoptera, including the red flour beetle *Tribolium castaneum*. Although this species lives in environments with minimal light, its behavior is rhythmic both under constant darkness and constant light, albeit not very robust (Reshma et al. 2024).

In the case of aphids and phylloxera, JET has been lost, while dCRY is still present. It is unclear whether the loss of JET triggered the accumulation of changes in dTIM, further permitting the loss of several exons at the 3’ end of the gene, or if some changes are influenced by coevolution between dTIM and PER proteins. Nevertheless, the major truncation of the C-terminal part suggests that the interaction between dTIM and dCRY is likely affected. Indeed, immunohistochemical localization of dCRY revealed that dCRY in the pea aphid, *A. pisum*, is stable upon light exposure (Collizi et al. 2021). Particularly interesting is the gradual reduction of clock components in *Blattodea*, where the loss of the dCRY gene is further accompanied by the loss of dTIM in termites (Kotwica-Rolinska et al. 2022a).

There are several lineages for which transcriptomic data suggest possible gene loss; however, well-assembled genomes, which are necessary to confirm such claims, are not available. For example, dCRY has not been identified in any representatives of Plecoptera or Raphidioptera. It is also important to note that some groups of organisms are insufficiently represented in GenBank, which hampers meaningful reconstruction of changes in the clock setup. For instance, Amphipoda, a group of Crustacea represented by *Parhyale hawaiensis*, lacks both dCRY and dTIM (Kwiatkowski et al. 2023). *Eurydice pulchra*, the closest species with dCRY and dTIM, diverged from amphipods more than 350 million years ago—around the same time when the ancestors of humans and *Xenopus* separated (Figure 8 and references used to refine the tree).

What are the potential consequences and selective forces driving the observed variability in the negative loop components of the circadian clock? Changes and losses affecting dTIM, dCRY, and JET are expected to influence the sensitivity of clocks to light, although parallel light input pathways involving opsins are also utilized (Helfrich-Forster et al. 2001). Reduced sensitivity to light might confer advantages under long photoperiods; however, the entrainment capacities of the clock may also depend on the neuroanatomy of the system, as demonstrated in flies from northern latitudes (Bertolini et al. 2019). Therefore, conducting functional analyses in specific organisms is essential to uncover the exact importance of particular genes.

It is important to note that the free-running period is a valuable marker of circadian clock properties. However, the actual selection pressure may act on adjusting the daily preference for locomotor activity. Clock genes also play a role in photoperiodic time measurement. Therefore, in cases such as geographic variability in the free-running period, it can be challenging to disentangle the effects of selection pressure on daily activity profiles from potential selection pressures acting on seasonal timing mechanisms (Pivarciova et al., 2016; Paolucci et al., 2019). Both dCRY+ and dCRY- insects exhibit robust photoperiodic timers (Stehlik et al. 2008; Ikeno et al. 2010; Urbanova et al. 2016; Kotwica-Rolinska et al. 2022b; Tobita and Kiuchi 2022). The interplay between daily anticipatory mechanisms and seasonal responses introduces additional selection pressures on circadian clock genes. Moreover, it is crucial to highlight that some circadian clock genes are also involved in regulating rhythms beyond the 24-hour range, such as the ∼13-hour tidal clock and circalunar (monthly) clocks (Kwiatkowski et al. 2023; Zantke et al. 2013; Zhang et al. 2023; Takekata et al. 2014).

## Material and Methods

For more detailed information on methodical description see supplementary methods, Supplementary Material online.

## Gene Identification and Phylogenetic Analyses

A systematic search for clock components was performed as was previously done to explore evolution, gene duplication, and gene losses of circadian clock components (Thakkar et al. 2022; Kotwica-Rolinska et al. 2022a) using protein and genomic databases, and transcriptome shotgun assemblies (TSA) in GenBank (NCBI). For details see supplementary methods, Supplementary Material online.

## Gene Absence and Gene Loss

In principle, gene loss becomes a parsimonious explanation if a particular gene is not found in entire monophyletic lineage for which a substantial amount of well-sequenced genomes and transcriptomes are available. See Table S1, supporting gene loss as a plausible explanation in specific cases.

## Prediction of Protein Domains

*D. melanogaster* dTIM protein domains presented by Lin et al. (2023) or identified in cell-based experiments (Saez and Young 1996) served as a reference for pairwise comparison with selected dTIMs. Nuclear localization signal (NLS) domains and acidic regions were predicted for each protein sequence *de novo*, see the supplementary methods. To calculate the length of the variable region in the central part of the protein and the C-terminal variable region (both depicted in Figure 2), conserved motifs were identified: YKDQ, located at the end of the dCRY-interaction domain; LLLR, located at the beginning of PER-binding site #2; DLIE, located in the second half of PER-binding site #2 sequence. The distances between specific sites were identified in Geneious Prime 2024 software (Biomatters, Auckland, New Zealand). The values were plotted as a dot for each sequence distance in Prism 7 (GraphPad Software, La Jolla, CA, USA) for every protein of our dataset.

## Substitutions per Amino Acid per Million Years

Phylogenetic trees were inferred from complete protein sequences in RAxML when the topology was forced to match the relatedness of organisms. The branch lengths from a common ancestor of Protostomia and Deuterostomia were determined for each protein, divided by 700 (the estimated separation of these lineages in millions of years), and plotted as individual dots using Prism 7 (GraphPad Software, La Jolla, CA, USA). The value thus corresponds to substitution per each amino acid position per million years. For details see the supplementary methods

## Gene Models and Exon Homologies

The gene models were either directly downloaded from GenBank species’s Whole-genome shotgun contigs (wgs) or Representative genomes (RefSeq genomes). Alternatively, gene models were manually reconstructed using genomic and transcriptomic data from GenBank, following two distinct scenarios: (1.) Long genomic contigs without annotated *timeless* genes. The wgs contigs were manually annotated by mapping Transcriptome Shotgun Assembly (TSA) and/or non-redundant nucleotide (nr/nt) *tim* sequences by the Minimap2 mapper (Li 2021) within Geneious Prime 2024 software (Biomatters, Auckland, New Zealand). Some gene models were reconstructed by mapping TSA sequences to an unannotated genomic scaffold. In the case of *Lepisma saccharina*, *Thermobia domestica d-tim* TSAs were used to scaffold and annotate DNA contigs. See tables S5 and S6 for accession numbers used to infer gene models.

In brief, similarities in gene structure were assessed as previously (Smykal et al. 2020) when conserved exon boundaries were depicted. In addition, similarity between exons was determined at the protein level (scoring matrix: BLOSUM90 with threshold 1) and depicted in a semi-quantitative scale with five similarity ranges (30-40; 41-50, 51-60, 61-70; 71-100 %). For details see the supplementary methods.

## Gene synteny

To reconstruct gene syntenies for *d-cry* and *jet* regions, we used a similar approach as was used previously (Smykal and Dolezel 2023). For details and for accession numbers, see the supplementary methods and Tables S3. and S4.

## Acknowledgments

We thank Martina Hajdušková (www.biographix.cz) for the color schemes used in Figure 8 and phylopic (https://www.phylopic.org/images) for schemes of starfish, sea urchin, and sea cucumber.

## Author Contributions

D.D. designed and guided the study, E.B. performed the protein comparisons, I.F. performed the phylogenetic analyses, P.C. reconstructed gene models under the supervision of V.S. who also performed synteny analyses. D.D. wrote the article and finalized the figures with input from all coauthors. Authors declare that they have no competing interests.

## References

1. Abrieux A, Xue Y, Cai Y, Lewald KM, Nguyen HN, Zhang Y, Chiu JC. 2020. EYES ABSENT and TIMELESS integrate photoperiodic and temperature cues to regulate seasonal physiology in Drosophila. Proc Natl Acad Sci U S A 117:15293–15304.

2. Aguillon R, Rinsky M, Simon-Blecher N, Doniger T, Appelbaum L, Levy O. 2024. CLOCK evolved in cnidaria to synchronize internal rhythms with diel environmental cues. Elife 12.

3. Allada R, White NE, So WV, Hall JC, Rosbash M. 1998. A mutant Drosophila homolog of mammalian Clock disrupts circadian rhythms and transcription of period and timeless. Cell 93:791–804.

4. Baretic D, Jenkyn-Bedford M, Aria V, Cannone G, Skehel M, Yeeles JTP. 2020. Cryo-EM Structure of the Fork Protection Complex Bound to CMG at a Replication Fork. Molecular Cell 78:926–940 e913.

5. Barnes JW, Tischkau SA, Barnes JA, Mitchell JW, Burgoon PW, Hickok JR, Gillette MU. 2003. Requirement of mammalian Timeless for circadian rhythmicity. Science 302:439–442.

6. Barbera M, Collantes-Alegre JM, Martinez-Torres D. 2017. Characterisation, analysis of expression and localisation of circadian clock genes from the perspective of photoperiodism in the aphid Acyrthosiphon pisum. Insect Biochem Mol Biol 83:54–67.

7. Bazalova O, Dolezel D. 2017. Daily Activity of the Housefly, Musca domestica, Is Influenced by Temperature Independent of 3’ UTR period Gene Splicing. G3 (Bethesda) 7:2637–2649.

8. Bazalova O, Kvicalova M, Valkova T, Slaby P, Bartos P, Netusil R, Tomanova K, Braeunig P, Lee HJ, Sauman I, et al. 2016. Cryptochrome 2 mediates directional magnetoreception in cockroaches. Proc Natl Acad Sci U S A 113:1660–1665.

9. Benna C, Bonaccorsi S, Wulbeck C, Helfrich-Forster C, Gatti M, Kyriacou CP, Costa R, Sandrelli F. 2010. Drosophila timeless2 is required for chromosome stability and circadian photoreception. Current Biology 20:346–352.

10. Bernatowicz PP, Kotwica-Rolinska J, Joachimiak E, Sikora A, Polanska MA, Pijanowska J, Bebas P. 2016. Temporal Expression of the Clock Genes in the Water Flea Daphnia pulex (Crustacea: Cladocera). J Exp Zool A Ecol Genet Physiol 325:233–254.

11. Bertolini E, Schubert FK, Zanini D, Sehadova H, Helfrich-Forster C, Menegazzi P. 2019. Life at High Latitudes Does Not Require Circadian Behavioral Rhythmicity under Constant Darkness. Current Biology 29:3928–3936 e3923.

12. Bhadra U, Thakkar N, Das P, Pal Bhadra M. 2017 Evolution of circadian rhythms: from bacteria to human. Sleep Med 35: 49–61.

13. Busza A, Emery-Le M, Rosbash M, Emery P. 2004. Roles of the two Drosophila CRYPTOCHROME structural domains in circadian photoreception. Science 304:1503–1506.

14. Cai YD, Chiu JC. 2021. Timeless in animal circadian clocks and beyond. Febs Journal. Febs Journal 289: 6559–6575.

15. Cai YD, Xue Y, Truong CC, Del Carmen-Li J, Ochoa C, Vanselow JT, Murphy KA, Li YH, Liu X, Kunimoto BL, et al. 2021. CK2 Inhibits TIMELESS Nuclear Export and Modulates CLOCK Transcriptional Activity to Regulate Circadian Rhythms. Current Biology 31:502–514 e507.

16. Ceriani MF, Darlington TK, Staknis D, Mas P, Petti AA, Weitz CJ, Kay SA. 1999. Light-dependent sequestration of TIMELESS by CRYPTOCHROME. Science 285:553–556.

17. Colizzi FS, Beer K, Cuti P, Deppisch P, Martinez Torres D, Yoshii T, Helfrich-Forster C. 2021. Antibodies Against the Clock Proteins Period and Cryptochrome Reveal the Neuronal Organization of the Circadian Clock in the Pea Aphid. Front Physiol 12:705048.

18. Collins B, Mazzoni EO, Stanewsky R, Blau J. 2006. Drosophila CRYPTOCHROME is a circadian transcriptional repressor. Current Biology 16:441–449.

19. Czarna A, Berndt A, Singh HR, Grudziecki A, Ladurner AG, Timinszky G, Kramer A, Wolf E. 2013. Structures of Drosophila cryptochrome and mouse cryptochrome1 provide insight into circadian function. Cell 153:1394–1405.

20. Danbara Y, Sakamoto T, Uryu O, Tomioka K. 2010. RNA interference of timeless gene does not disrupt circadian locomotor rhythms in the cricket Gryllus bimaculatus. Journal of Insect Physiology 56:1738–1745.

21. D’Costa A, Reifegerste R, Sierra S, Moses K. 2006. The Drosophila ramshackle gene encodes a chromatin-associated protein required for cell morphology in the developing eye. Mech Dev 123:591–604.

22. Deppisch P, Prutscher JM, Pegoraro M, Tauber E, Wegener C, Helfrich-Forster C. 2022. Adaptation of Drosophila melanogaster to Long Photoperiods of High-Latitude Summers Is Facilitated by the ls-Timeless Allele. J Biol Rhythms:7487304221082448.

23. Deppisch P, Kirsch V, Helfrich-Forster C, Senthilan PR. 2023. Contribution of cryptochromes and photolyases for insect life under sunlight. J Comp Physiol A Neuroethol Sens Neural Behav Physiol.

24. Dolezel D. 2023. Molecular Mechanism of the Circadian Clock. In: Numata H, Tomioka K (ed) Insect Chronobiology, 1st edn. Springer, Singapore; ISBN 978-981-99-0725-0. 49-84.

25. Dolezelova E, Dolezel D, Hall JC. 2007. Rhythm defects caused by newly engineered null mutations in Drosophila’s cryptochrome gene. Genetics 177:329–345.

26. Edgar RS, Green EW, Zhao Y, van Ooijen G, Olmedo M, Qin X, Xu Y, Pan M, Valekunja UK, Feeney KA, et al. 2012. Peroxiredoxins are conserved markers of circadian rhythms. Nature 485:459–464.

27. Emery P, So WV, Kaneko M, Hall JC, Rosbash M. 1998. CRY, a Drosophila clock and light-regulated cryptochrome, is a major contributor to circadian rhythm resetting and photosensitivity. Cell 95:669–679.

28. Emery P, Stanewsky R, Hall JC, Rosbash M. 2000a. Drosophila cryptochromes - A unique circadian-rhythm photoreceptor. Nature 404:456–457.

29. Emery P, Stanewsky R, Helfrich-Forster C, Emery-Le M, Hall JC, Rosbash M. 2000b. Drosophila CRY is a deep brain circadian photoreceptor. Neuron 26:493–504.

30. Foley LE, Ling J, Joshi R, Evantal N, Kadener S, Emery P. 2019. Drosophila PSI controls circadian period and the phase of circadian behavior under temperature cycle via tim splicing. Elife 8.

31. Giesecke A, Johnstone PS, Lamaze A, Landskron J, Atay E, Chen KF, Wolf E, Top D, Stanewsky R. 2023. A novel period mutation implicating nuclear export in temperature compensation of the Drosophila circadian clock. Current Biology 33:336–350 e335.

32. Gotter AL, Manganaro T, Weaver DR, Kolakowski LF, Possidente B, Sriram S, MacLaughlin DT, Reppert SM. 2000. A time-less function for mouse Timeless. Nature Neuroscience 3:755–756.

33. Grabarczyk DB. 2020. Crystal structure and interactions of the Tof1-Csm3 (Timeless-Tipin) fork protection complex. Nucleic Acids Research 48:6996–7004.

34. Hasegawa K, Saigusa T, Tamai Y. 2005. Caenorhabditis elegans opens up new insights into circadian clock mechanisms. Chronobiology International 22:1–19.

35. Helfrich-Forster C, Winter C, Hofbauer A, Hall JC, Stanewsky R. 2001. The circadian clock of fruit flies is blind after elimination of all known photoreceptors. Neuron 30:249–261.

36. Iiams SE, Wan G, Zhang J, Lugena AB, Zhang Y, Hayden AN, Merlin C. 2024. Loss of functional cryptochrome 1 reduces robustness of 24-hour behavioral rhythms in monarch butterflies. iScience 27:108980.

37. Ikeno T, Tanaka SI, Numata H, Goto SG. 2010. Photoperiodic diapause under the control of circadian clock genes in an insect. Bmc Biology 8.

38. Jang AR, Moravcevic K, Saez L, Young MW, Sehgal A. 2015. Drosophila TIM Binds Importin alpha1, and Acts as an Adapter to Transport PER to the Nucleus. Plos Genetics 11:e1004974.

39. Kamae Y, Tomioka K. 2012. timeless is an essential component of the circadian clock in a primitive insect, the firebrat Thermobia domestica. J Biol Rhythms 27:126–134.

40. Kaniewska MM, Vaneckova H, Dolezel D, Kotwica-Rolinska J. 2020. Light and Temperature Synchronizes Locomotor Activity in the Linden Bug, Pyrrhocoris apterus. Front Physiol 11:242.

41. Kobelkova A, Bajgar A, Dolezel D. 2010. Functional Molecular Analysis of a Circadian Clock Gene timeless Promoter from the Drosophilid Fly Chymomyza costata. Journal of Biological Rhythms 25:399–409.

42. Koh K, Zheng X, Sehgal A. 2006. JETLAG resets the Drosophila circadian clock by promoting light-induced degradation of TIMELESS. Science 312:1809–1812.

43. Koike N, Hida A, Numano R, Hirose M, Sakaki Y, Tei H. 1998. Identification of the mammalian homologues of the Drosophila timeless gene, Timeless1. Febs Letters 441:427-431.

44. Kostal V. 2006. Eco-physiological phases of insect diapause. Journal of Insect Physiology 52:113–127.

45. Kotwica-Rolinska J, Chodakova L, Smykal V, Damulewicz M, Provaznik J, Wu BC, Hejnikova M, Chvalova D, Dolezel D. 2022. Loss of Timeless Underlies an Evolutionary Transition within the Circadian Clock. Molecular Biology and Evolution 39.

46. Kotwica-Rolinska J, Damulewicz M, Chodakova L, Kristofova L, Dolezel D. 2022. Pigment Dispersing Factor Is a Circadian Clock Output and Regulates Photoperiodic Response in the Linden Bug, Pyrrhocoris apterus. Front Physiol 13:884909.

47. Kume K, Zylka MJ, Sriram S, Shearman LP, Weaver DR, Jin XW, Maywood ES, Hastings MH, Reppert SM. 1999. mCRY1 and mCRY2 are essential components of the negative limb of the circadian clock feedback loop. Cell 98:193–205.

48. Kurien P, Hsu PK, Leon J, Wu D, McMahon T, Shi G, Xu Y, Lipzen A, Pennacchio LA, Jones CR, et al. 2019. TIMELESS mutation alters phase responsiveness and causes advanced sleep phase. Proc Natl Acad Sci U S A 116:12045–12053.

49. Kwiatkowski ER, Schnytzer Y, Rosenthal JJC, Emery P. 2023. Behavioral circatidal rhythms require Bmal1 in Parhyale hawaiensis. Current Biology 33:1867–1882 e1865.

50. Kwiatkowski ER, Emery P. 2024. Cnidarians are CLOCKing in. Elife 13.

51. Lamaze A, Chen C, Leleux S, Xu M, George R, Stanewsky R. 2022. A natural timeless polymorphism allowing circadian clock synchronization in "white nights". Nature Communications 13:1724.

52. Levy C, Zoltowski BD, Jones AR, Vaidya AT, Top D, Widom J, Young MW, Scrutton NS, Crane BR, Leys D. 2013. Updated structure of Drosophila cryptochrome. Nature 495:E3–4.

53. Li H. 2021. New strategies to improve minimap2 alignment accuracy. Bioinformatics 37(23): 4572–4574.

54. Lin C, Feng S, DeOliveira CC, Crane BR. 2023. Cryptochrome-Timeless structure reveals circadian clock timing mechanisms. Nature 617:194–199.

55. Lindestad O, Nylin S, Wheat CW, Gotthard K. 2022. Local adaptation of life cycles in a butterfly is associated with variation in several circadian clock genes. Molecular Ecology 31:1461–1475.

56. Martin Anduaga A, Evantal N, Patop IL, Bartok O, Weiss R, Kadener S. 2019. Thermosensitive alternative splicing senses and mediates temperature adaptation in Drosophila. Elife 8.

57. Matsumoto A, Tomioka K, Chiba Y, Tanimura T. 1999. timrit Lengthens circadian period in a temperature-dependent manner through suppression of PERIOD protein cycling and nuclear localization. Molecular and Cellular Biology 19:4343–4354.

58. Mendoza-Viveros L, Bouchard-Cannon P, Hegazi S, Cheng AH, Pastore S, Cheng HM. 2017. Molecular modulators of the circadian clock: lessons from flies and mice. Cellular and Molecular Life Sciences. 74:1035–1059

59. Meyer P, Saez L, Young MW. 2006. PER-TIM interactions in living Drosophila cells: An interval timer for the circadian clock. Science 311:226–229.

60. Michael AK, Stoos L, Crosby P, Eggers N, Nie XY, Makasheva K, Minnich M, Healy KL, Weiss J, Kempf G, et al. 2023. Cooperation between bHLH transcription factors and histones for DNA access. Nature 619:385–393.

61. Montelli S, Mazzotta G, Vanin S, Caccin L, Corra S, De Pitta C, Boothroyd C, Green EW, Kyriacou CP, Costa R. 2015. period and timeless mRNA Splicing Profiles under Natural Conditions in Drosophila melanogaster. J Biol Rhythms 30:217–227.

62. Nose M, Tokuoka A, Bando T, Tomioka K. 2017. timeless2 plays an important role in reproduction and circadian rhythms in the cricket Gryllus bimaculatus. Journal of Insect Physiology. 105:9–17

63. Ozturk N, Vanvickle-Chavez SJ, Akileswaran L, Van Gelder RN, Sancar A. 2013. Ramshackle (Brwd3) promotes light-induced ubiquitylation of Drosophila Cryptochrome by DDB1-CUL4-ROC1 E3 ligase complex. Proc Natl Acad Sci U S A. 110: 4980–4985.

64. Paolucci, S., Dalla Benetta, E., Salis, L., Dolezel, D., van de Zande, L., Beukeboom, L.W., 2019. Latitudinal Variation in Circadian Rhythmicity in Nasonia vitripennis. Behavioral sciences 9.

65. Peschel N, Chen KF, Szabo G, Stanewsky R. 2009. Light-Dependent Interactions between the Drosophila Circadian Clock Factors Cryptochrome, Jetlag, and Timeless. Current Biology 19:241–247.

66. Peschel N, Veleri S, Stanewsky R. 2006. Veela defines a molecular link between Cryptochrome and Timeless in the light-input pathway to Drosophila’s circadian clock. Proceedings of the National Academy of Sciences of the United States of America 103:17313–17318.

67. Pivarciova L, Vaneckova H, Provaznik J, Wu BC, Pivarci M, et al. 2016. Unexpected Geographic Variability of the Free Running Period in the Linden Bug *Pyrrhocoris apterus*. Journal of Biological Rhythms 31:568–576

68. Poupardin R, Schottner K, Korbelova J, Provaznik J, Dolezel D, Pavlinic D, Benes V, Kostal V. 2015. Early transcriptional events linked to induction of diapause revealed by RNAseq in larvae of drosophilid fly, Chymomyza costata. Bmc Genomics 16:720.

69. Putker M, Wong DCS, Seinkmane E, Rzechorzek NM, Zeng A, Hoyle NP, Chesham JE, Edwards MD, Feeney KA, Fischer R, et al. 2021. CRYPTOCHROMES confer robustness, not rhythmicity, to circadian timekeeping. Embo Journal:e106745.

70. Reshma R, Pruser T, Schulz NKE, Mayer PMF, Ogueta M, Stanewsky R, Kurtz J. 2024. Deciphering a Beetle Clock: Individual and Sex-Dependent Variation in Daily Activity Patterns. J Biol Rhythms:7487304241263619.

71. Rosato E, Trevisan A, Sandrelli F, Zordan M, Kyriacou CP, Costa R. 1997. Conceptual translation of timeless reveals alternative initiating methionines in Drosophila. Nucleic Acids Research 25:455–458.

72. Rothenfluh A, Abodeely M, Price JL, Young MW. 2000a. Isolation and analysis of six timeless alleles that cause short- or long-period circadian rhythms in Drosophila. Genetics 156:665–675.

73. Rothenfluh A, Young MW, Saez L. 2000b. A TIMELESS-independent function for PERIOD proteins in the Drosophila clock. Neuron 26:505–514.

74. Rubin EB, Shemesh Y, Cohen M, Elgavish S, Robertson HM, Bloch G. 2006. Molecular and phylogenetic analyses reveal mammalian-like clockwork in the honey bee (Apis mellifera) and shed new light on the molecular evolution of the circadian clock. Genome Research 16:1352–1365.

75. Rutila JE, Suri V, Le M, So WV, Rosbash M, Hall JC. 1998. CYCLE is a second bHLH-PAS clock protein essential for circadian rhythmicity and transcription of Drosophila period and timeless. Cell 93:805–814.

76. Saez L, Young MW. 1996. Regulation of nuclear entry of the Drosophila clock proteins period and timeless. Neuron 17:911–920.

77. Saez L, Derasmo M, Meyer P, Stieglitz J, Young MW. 2011. A Key Temporal Delay in the Circadian Cycle of Drosophila Is Mediated by a Nuclear Localization Signal in the Timeless Protein. Genetics 188:591–U166.

78. Sandrelli F, Tauber E, Pegoraro M, Mazzotta G, Cisotto P, Landskron J, Stanewsky R, Piccin A, Rosato E, Zordan M, et al. 2007. A molecular basis for natural selection at the timeless locus in Drosophila melanogaster. Science 316:1898–1900.

79. Saunders DS, Henrich VC, Gilbert LI. 1989. Induction of diapause in Drosophila melanogaster: photoperiodic regulation and the impact of arrhythmic clock mutations on time measurement. Proc Natl Acad Sci U S A 86:3748–3752.

80. Shakhmantsir I, Nayak S, Grant GR, Sehgal A. 2018. Spliceosome factors target timeless (tim) mRNA to control clock protein accumulation and circadian behavior in Drosophila. Elife 7.

81. Singh S, Giesecke A, Damulewicz M, Fexova S, Mazzotta GM, Stanewsky R, Dolezel D. 2019. New Drosophila Circadian Clock Mutants Affecting Temperature Compensation Induced by Targeted Mutagenesis of Timeless. Front Physiol 10:1442.

82. Smýkal V, Pivarči M, Provazník J, Bazalová O, Jedlička P, Lukšan O, Horák A, Vaněčková H, Beneš V, Fiala I, et al. 2020. Complex evolution of insect insulin receptors and homologous decoy receptors, and functional significance of their multiplicity. Mol Biol Evol. 37(6):1775–1789.

83. Smykal V, Dolezel D. 2023. Evolution of proteins involved in the final steps of juvenile hormone synthesis. Journal of Insect Physiology 145:104487.

84. Stanewsky R, Kaneko M, Emery P, Beretta B, Wager-Smith K, Kay SA, Rosbash M, Hall JC. 1998. The cry(b) mutation identifies cryptochrome as a circadian photoreceptor in Drosophila. Cell 95:681–692.

85. Stehlik J, Zavodska R, Shimada K, Sauman I, Kostal V. 2008. Photoperiodic induction of diapause requires regulated transcription of timeless in the larval brain of Chymomyza costata. Journal of Biological Rhythms 23:129–139.

86. Takekata H, Numata H, Shiga S, Goto SG. 2014. Silencing the circadian clock gene Clock using RNAi reveals dissociation of the circatidal clock from the circadian clock in the mangrove cricket. Journal of Insect Physiology 68C:16-22.

87. Tauber E, Zordan M, Sandrelli F, Pegoraro M, Osterwalder N, Breda C, Daga A, Selmin A, Monger K, Benna C, et al. 2007. Natural selection favors a newly derived timeless allele in Drosophila melanogaster. Science 316:1895–1898.

88. Thakkar N, Giesecke A, Bazalova O, Martinek J, Smykal V, Stanewsky R, Dolezel D. 2022. Evolution of casein kinase 1 and functional analysis of new doubletime mutants in Drosophila. Front Physiol 13:1062632.

89. Thakkar N, Hejzlarova A, Brabec V, Dolezel D. 2023. Germline Editing of Drosophila Using CRISPR- Cas9-based Cytosine and Adenine Base Editors. CRISPR J.

90. Tobita H, Kiuchi T. 2024. Knockout of cryptochrome 1 disrupts circadian rhythm and photoperiodic diapause induction in the silkworm, Bombyx mori. Insect Biochem Mol Biol:104153.

91. Tobita H, Kiuchi T. 2022. Knockouts of positive and negative elements of the circadian clock disrupt photoperiodic diapause induction in the silkworm, Bombyx mori. Insect Biochem Mol Biol 149:103842.

92. Tomioka K, Matsumoto A. 2015. Circadian molecular clockworks in non-model insects. Current Opinion in Insect Science 7:58–64.

93. Top D, Harms E, Syed S, Adams EL, Saez L. 2016. GSK-3 and CK2 Kinases Converge on Timeless to Regulate the Master Clock. Cell Rep 16: 357–367.

94. Tumova S, Dolezel D, M. J. 2024. Conserved and Unique Roles of bHLH-PAS Transcription Factors in Insects – From Clock to Hormone Reception. Journal of Molecular Biology 436 (2024) 168332:1-25.

95. Unsal-Kacmaz K, Mullen TE, Kaufmann WK, Sancar A. 2005. Coupling of human circadian and cell cycles by the timeless protein. Molecular and Cellular Biology 25:3109–3116.

96. Urbanova V, Bazalova O, Vaneckova H, Dolezel D. 2016. Photoperiod regulates growth of male accessory glands through juvenile hormone signaling in the linden bug, Pyrrhocoris apterus. Insect Biochem Mol Biol 70:184–190.

97. Werckenthin A, Huber J, Arnold T, Koziarek S, Plath MJA, Plath JA, Stursberg O, Herzel H, Stengl M. 2020. Neither per, nor tim1, nor cry2 alone are essential components of the molecular circadian clockwork in the Madeira cockroach. Plos One 15:e0235930.

98. Yildiz O, Doi M, Yujnovsky I, Cardone L, Berndt A, Hennig S, Schulze S, Urbanke C, Sassone-Corsi P, Wolf E. 2005. Crystal structure and interactions of the PAS repeat region of the Drosophila clock protein PERIOD. Molecular Cell 17:69–82.

99. Yuan Q, Metterville D, Briscoe AD, Reppert SM. 2007. Insect cryptochromes: Gene duplication and loss define diverse ways to construct insect circadian clocks. Molecular Biology and Evolution 24:948–955.

100. Zantke J, Ishikawa-Fujiwara T, Arboleda E, Lohs C, Schipany K, Hallay N, Straw AD, Todo T, Tessmar-Raible K. 2013. Circadian and circalunar clock interactions in a marine annelid. Cell Rep 5:99–113.

101. Zhang L, Green EW, Webster SG, Hastings MH, Wilcockson DC, Kyriacou CP. 2023. Correction: The circadian clock gene bmal1 is necessary for co-ordinated circatidal rhythms in the marine isopod Eurydice pulchra (Leach). Plos Genetics 19:e1011047.

102. Zoltowski BD, Vaidya AT, Top D, Widom J, Young MW, Crane BR. 2011. Structure of full-length Drosophila cryptochrome. Nature 480:396–U156.

