## Supplementary material for "Coevolution of *Drosophila*-type Timeless with Partner Clock Proteins"

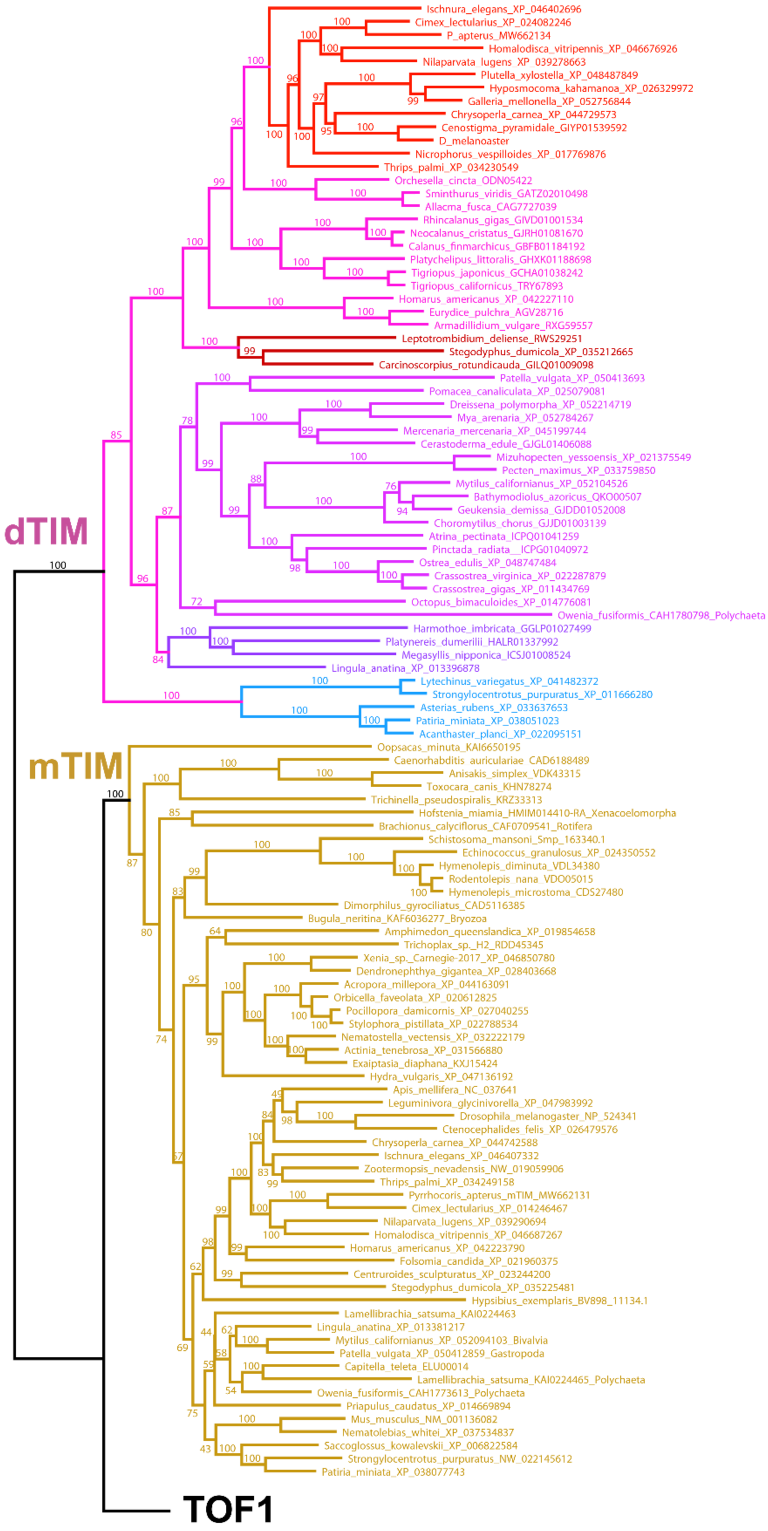


**Supplementary Fig. 1, Part A.** Phylogeny of dTIM (*Drosophila*-type timeless), mTIM (mammalian-type timeless), and TOF1 (Topoisomerase 1-associated Factor). The unrooted tree was inferred from protein sequence alignment using the RAxML maximum likelihood GAMMA-based model (final GAMMA-based score of the best tree is -328285.145114). Bootstrap values, inferred from 500 replicates, are displayed as percentages.

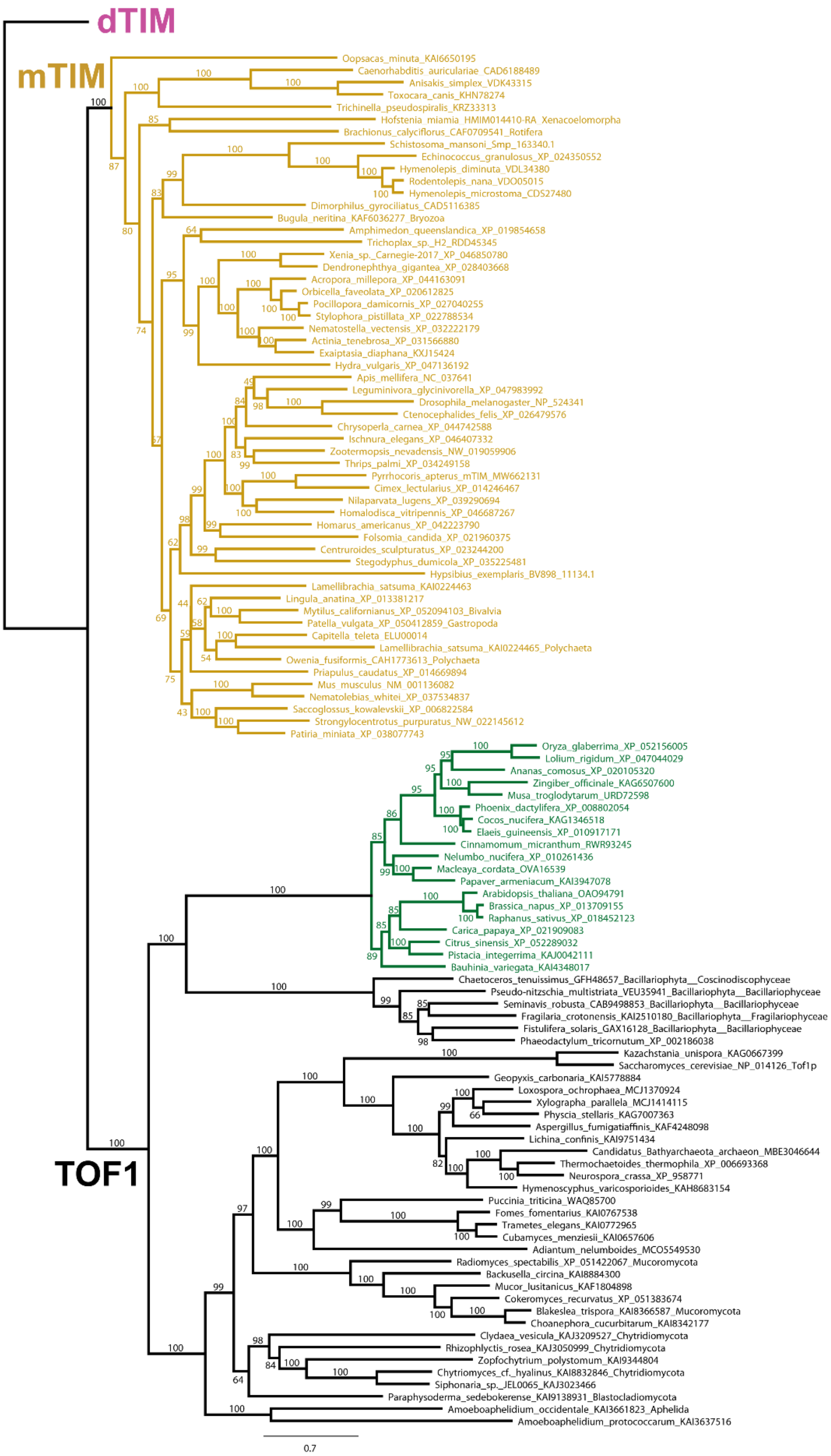


**Supplementary Fig. 1 partB.** Phylogeny of dTIM, mTIM (mammalian-type timeless), and TOF1. The unrooted tree inferred from protein sequence alignment using RAxML maximum likelihood GAMMA-based model (final GAMMA-based score of the best tree is -328285.145114) in Geneious Prime 2024.0.7. Bootstrap values, inferred from 500 replicates, are displayed as percentages.

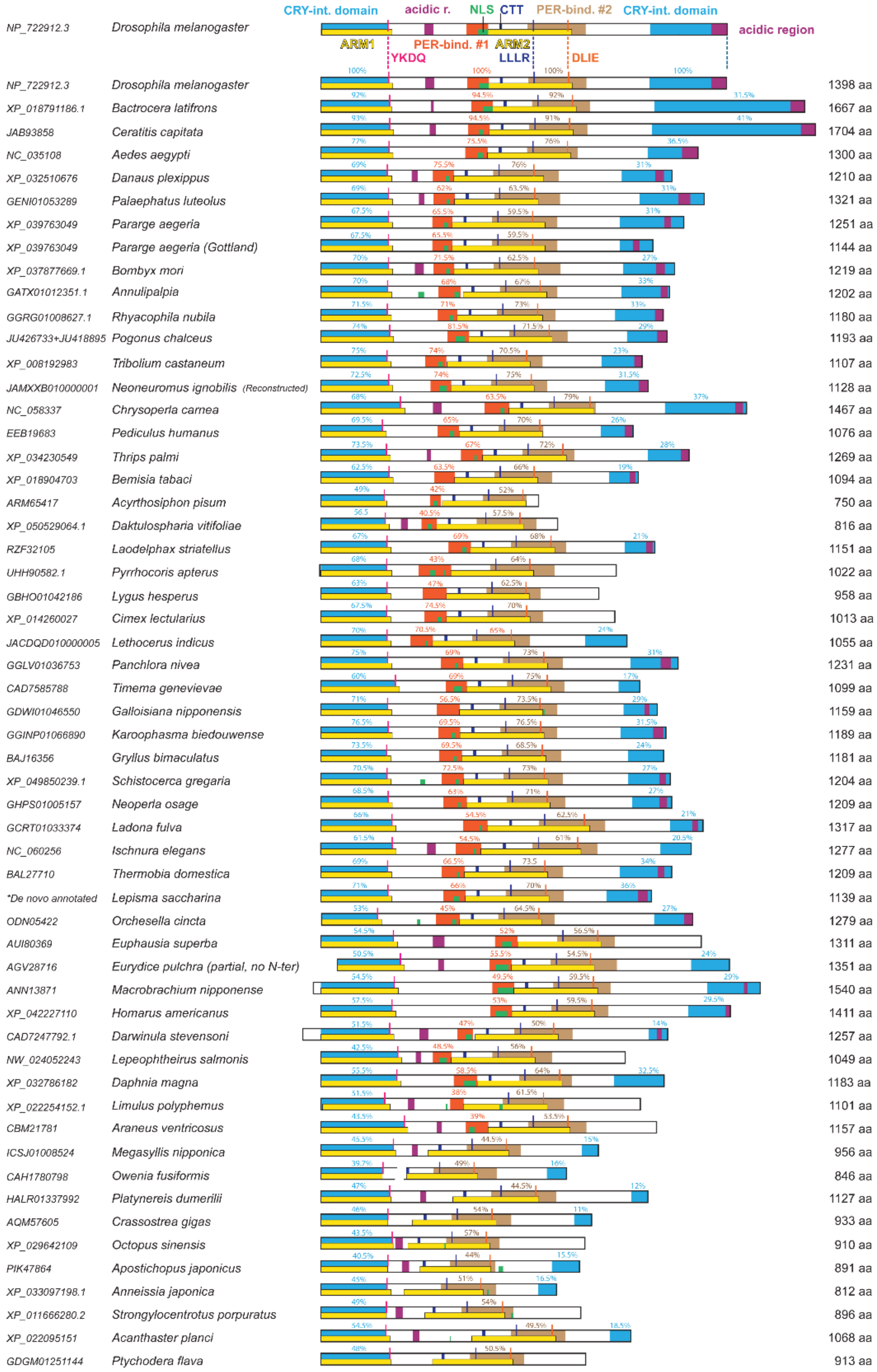


**Supplementary Fig. 2.** Protein models of representative dTIM sequences with highlighted sequence similarities corresponding to *Drosophila* dTIM domains. In the case of PER-bind site #1 and the CRY-interaction domain, even very low similarities are depicted, and the exact percentage values are shown. Note that the variability involves the inter-armadillo region (between ARM1 and ARM2) with PER-binding site #1 and the C-terminal part, whereas both armadillo regions (ARM1, ARM2) and PER-binding site #2 are generally conserved.

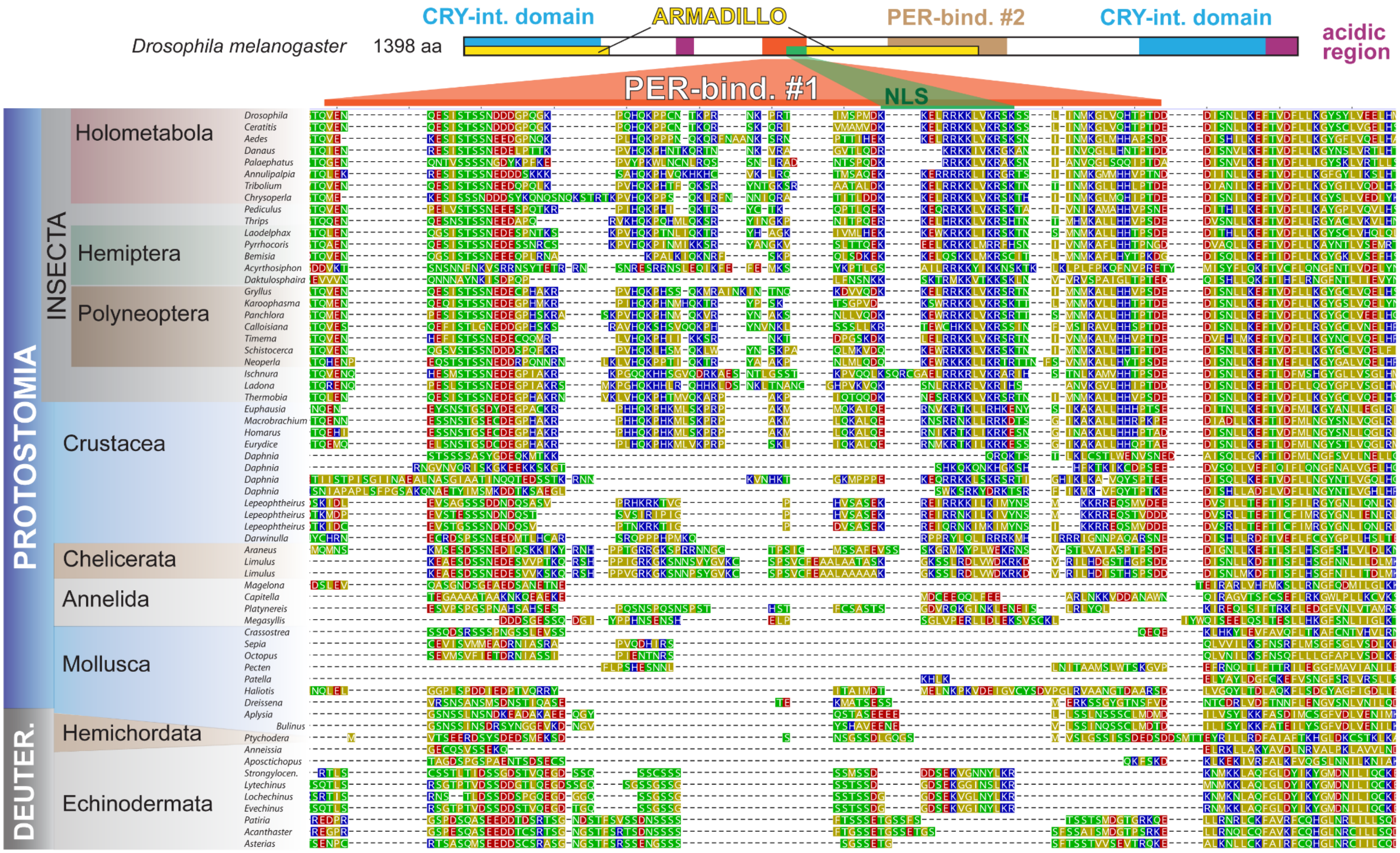


**Supplementary Fig. 3**. The alignment of PER-binding site #1 (functionally identified for *Drosophila* dTIM; Saez and Young 1996) indicates that this region is well-conserved in insect, Crustacea and contains nuclear localization signal (NLS). While the corresponding region can still be aligned in chelicerates, it differs significantly in annelids, mollusks, and deuterostomian species. Notably, substantial sequence diversity is observed among echinoderm species.

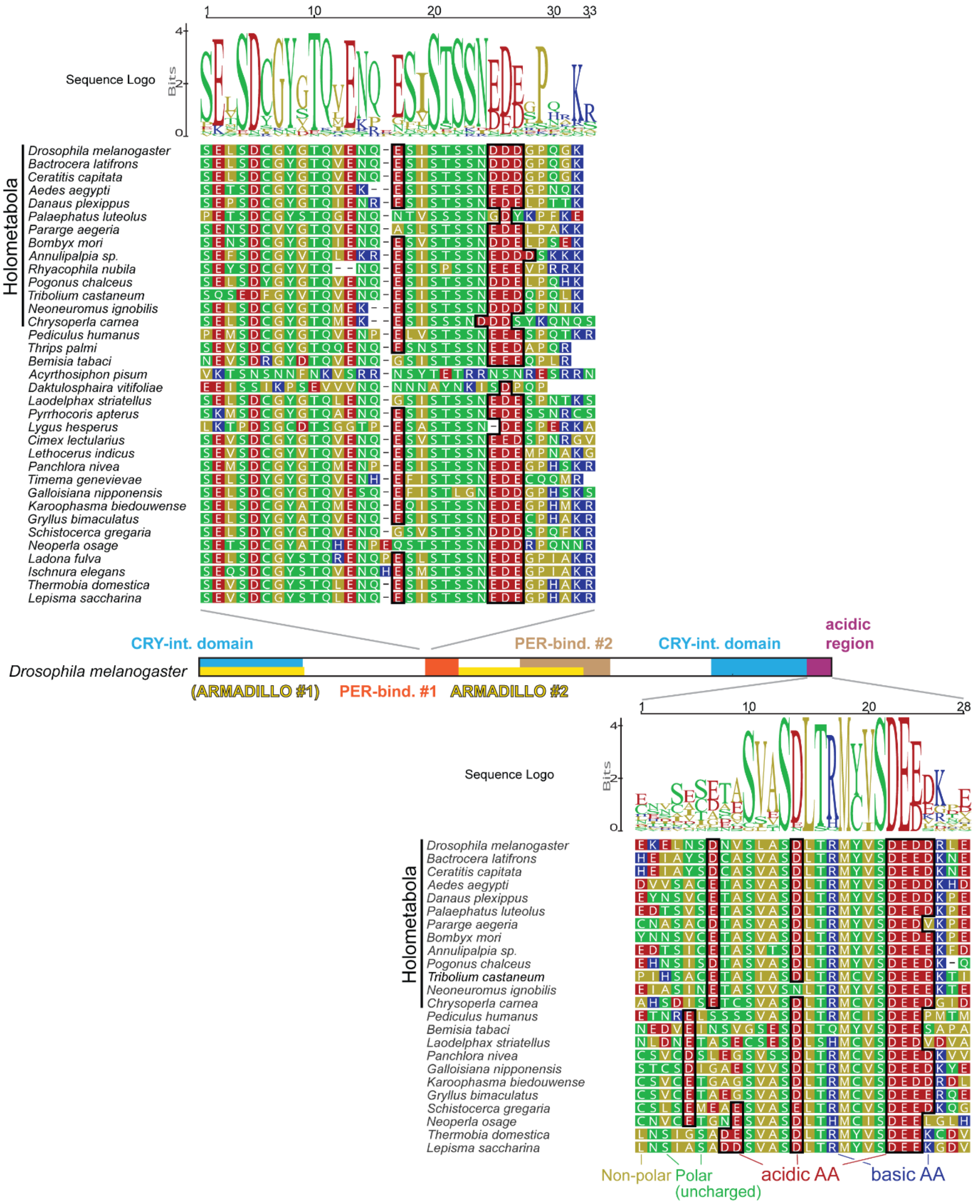


**Supplementary Fig. 4.** Two acidic regions are found in nearly all insect dTIMs. The top alignment showes the pattern located at the beginning of PER-binding site #1 (see schematic *Drosophila* dTIM in the middle of the page for key dTIM functional domains). This motif is found in all analyzed insects except phylloxera and aphids (represented by *Daktulosphaira* and *Acyrthosiphon*, respectively). The bottom alignment highlights the acidic region located at the C-tail of dTIM, which is present in most insects, except Odonata, Mantophasmatodea, and Grylloblattodea. In some hemipteran species, the C-tail motif has been completely lost (species without the motif were not shown). The sequences were aligned in Geneious Prime using MAFFT alignment.

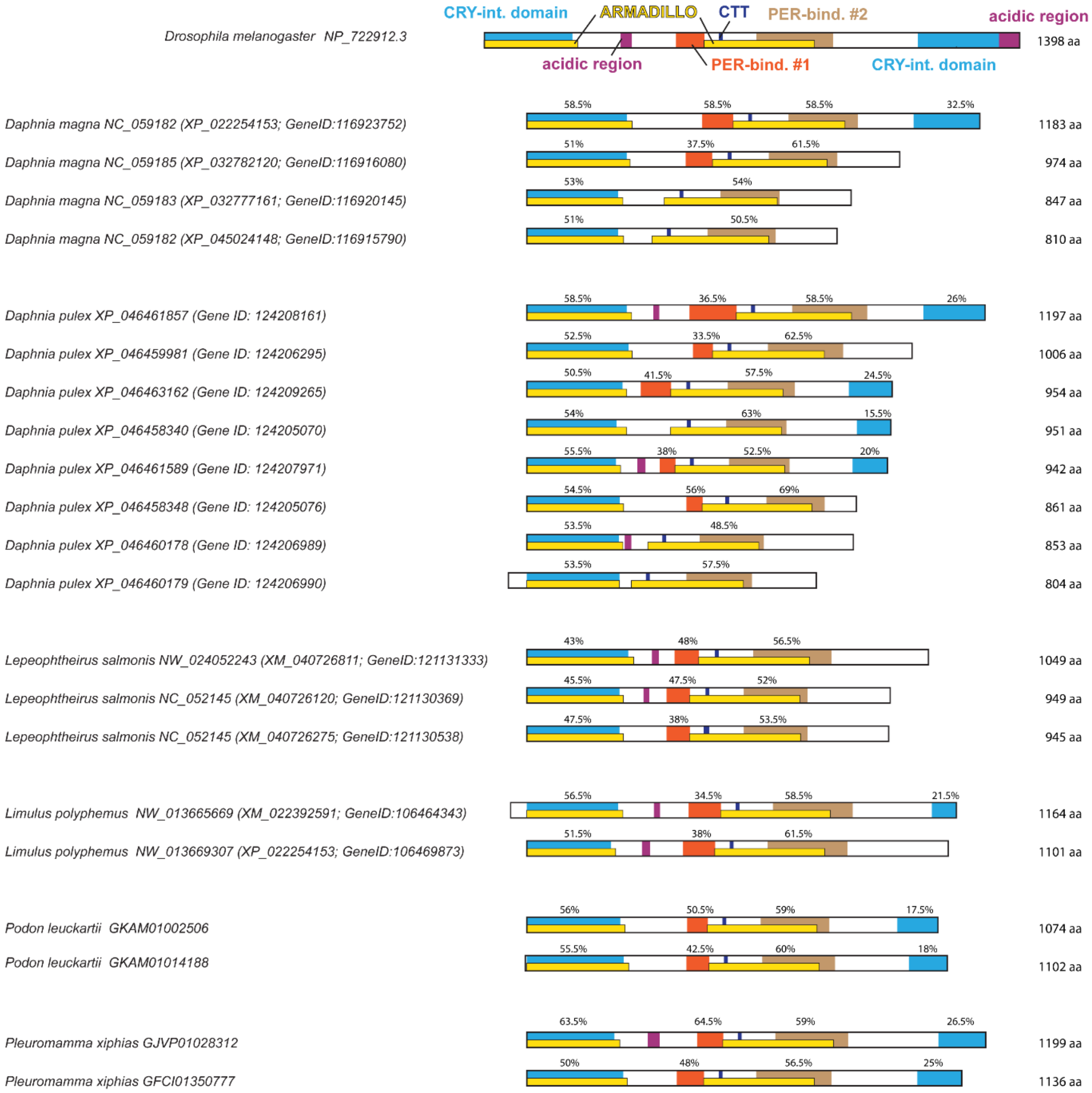


**Supplementary Fig. 5.** Detailed description of protein domains in species with multiple dTIM paralogs (extension of data presented in Figure 3). Protein domains were identified and color-highlighted based on sequence similarity to the corresponding domains in Drosophila dTIM (top). The armadillo domains (ARM1 and ARM2) and PER-binding site #2 are generally well-conserved, with similarity exceeding 40%. In contrast, significant variability is observed in the inter-armadillo region (between ARM1 and ARM2), which includes PER-binding site #1, as well as in the C-terminal region. Exact percentage similarity values are provided for the PER-binding site #1 and the CRY interaction domain.

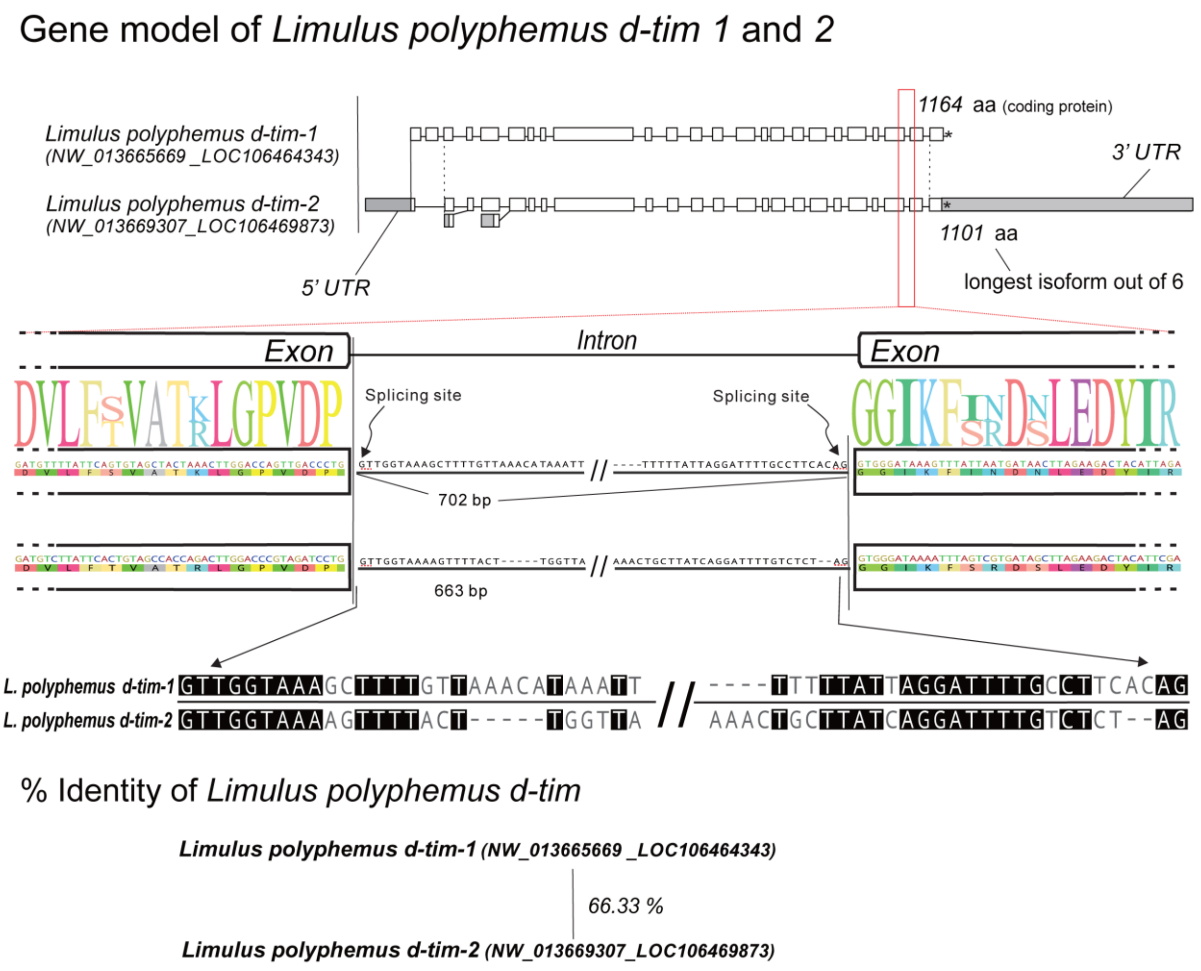


**Supplementary Fig. 6.** The gene model illustrates that both *Limulus polyphemus d-tim* paralogs share a comparable gene structure. The detail of the penultimate intron with surrounding exon sequences points to several sequence differences.

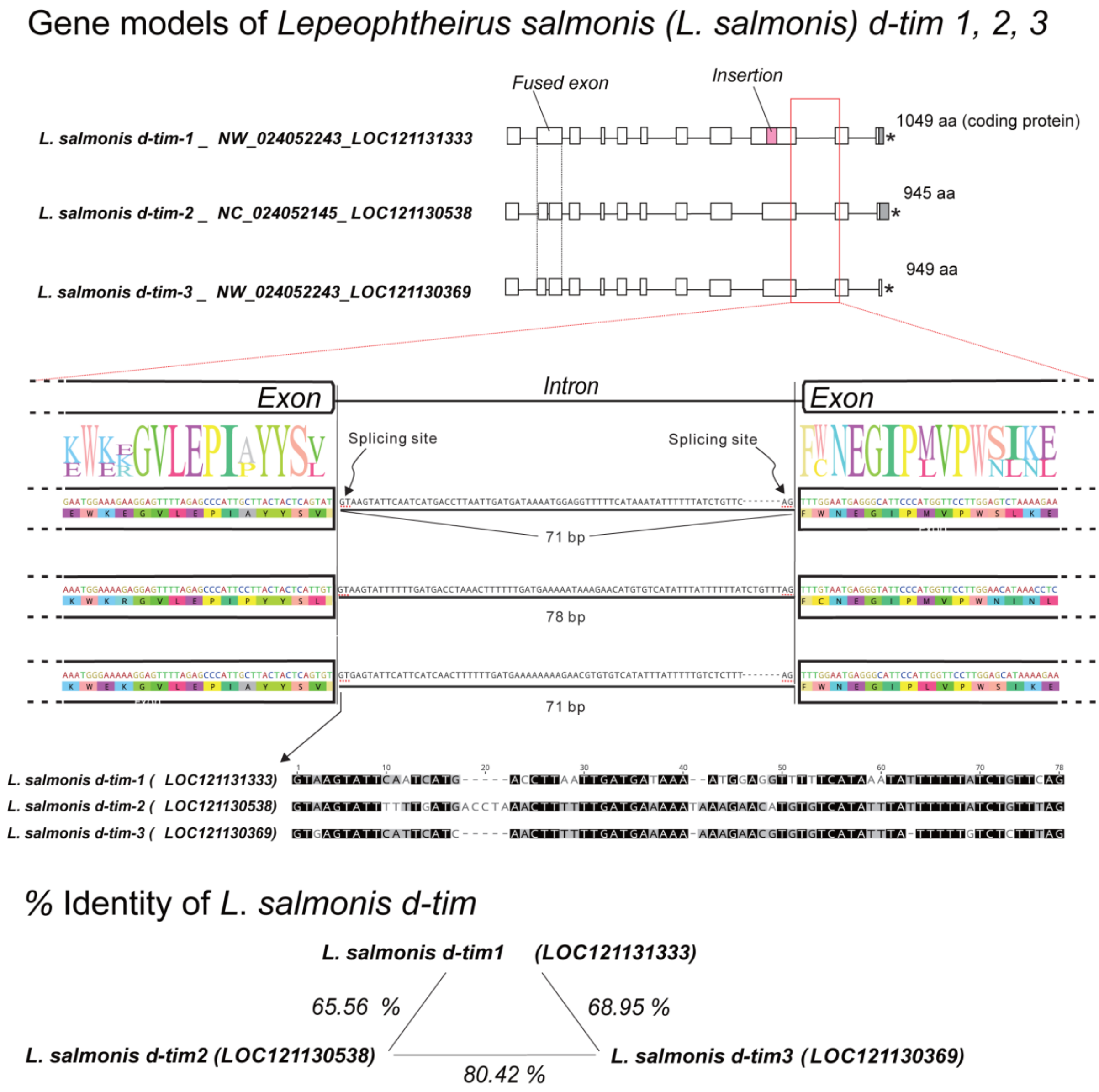


**Supplementary Fig. 7.** The gene model illustrates that all three *Lepeophteirus salmonis d-tim* paralogs share a comparable gene structure. The detail of the penultimate intron with surrounding exon sequences points to several sequence differences.

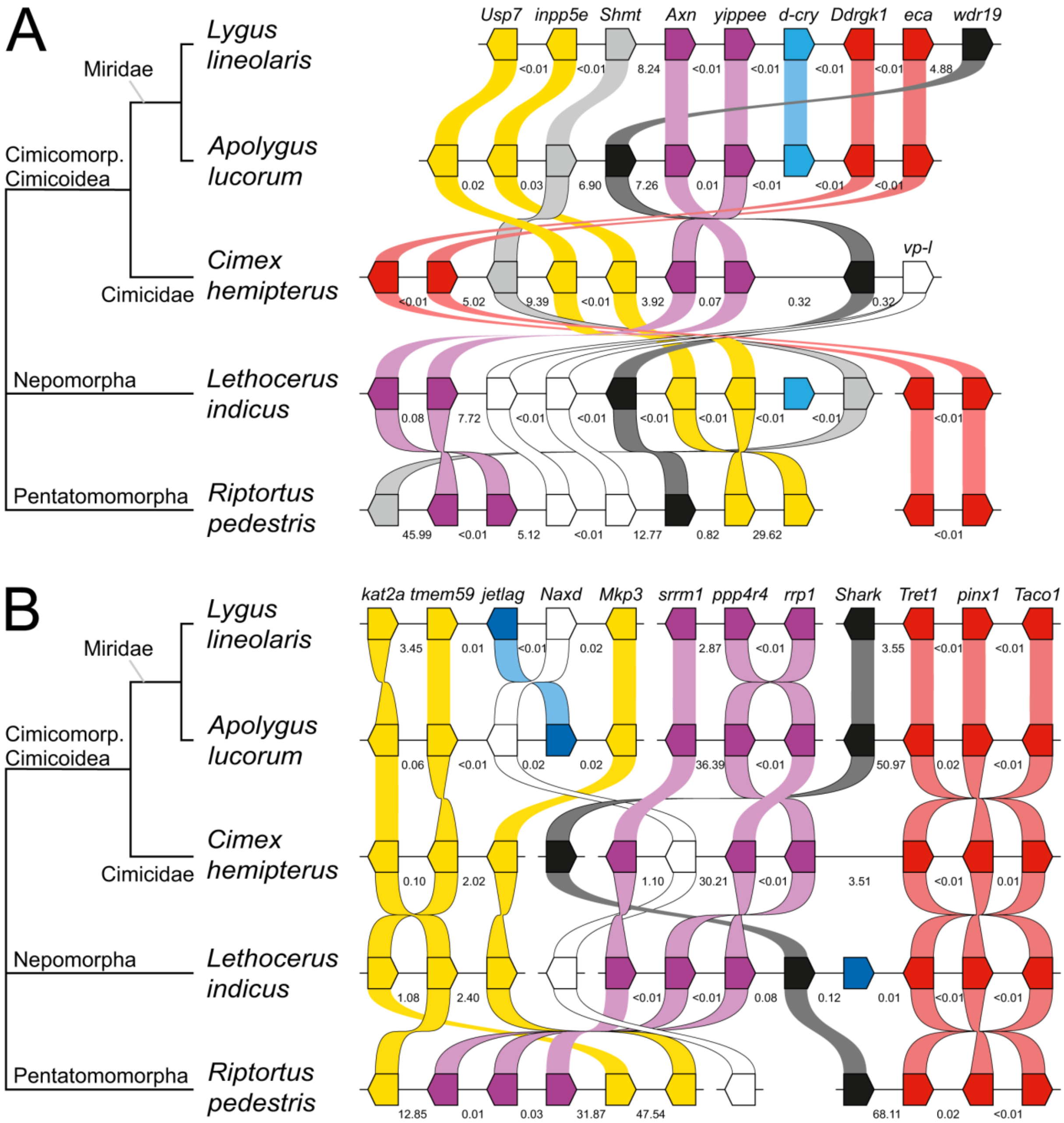


**Supplementary Fig. 8.** Extended depiction of gene syntenies presented in fig. 5. The gene synteny analysis shows the independent loss of *Drosophila*-type *cryptochrome* (*d-cry*) and *jetlag* (*jet*) genes in *Cimex hemipterus* and *Riptortus pedestris*. Panels show (A) *d-cry* and (B) *jet* gene synteny across five (pan)heteropteran species. The genomic localization of *d-cry* and *jet* in *L. lineolaris*/*A. lucorum* (Miridae) and *L. indicus* (Nepomorpha) differ, as indicated by distinct sets of protein-coding genes surrounding both loci. While syntenic genes can still be mapped to *C. hemipterus* (Cimicidae) and *R. pedestris* (Pentatomorpha) genomes, differences in gene composition and relative orientation support the independent loss of *d-cry* and *jet*. Horizontal black lines represent genomic contigs/scaffolds (not to scale), with arrow-like boxes denoting protein-coding genes and their orientations. Orthologous genes are color-coded and connected, and numbers between genes (arrow-like boxes) indicate intergenic distances in millions of base pairs (Mbp).

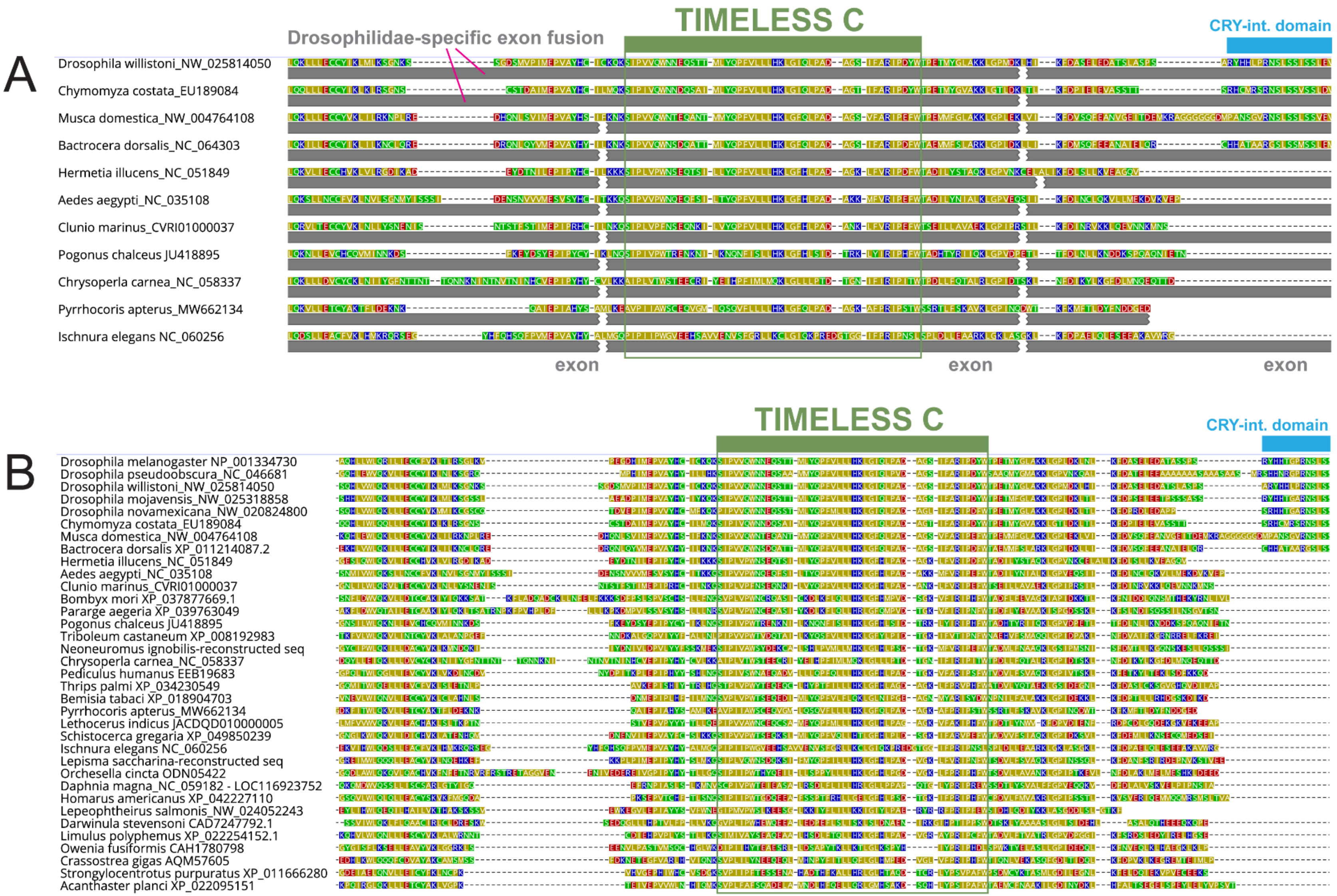


**Supplementary Fig. 9.** Details of the TIMELESS C domain. (A) Portion of insect dTIM sequences with annotated exons. Note the exon fusion in *Drosophila* and *Chymomyza*. (B) Detail of dTIM representing all major groups of Bilateria. TIMELESS C domain is highlighted by the green bar and rectangle.

### Supplementary Material and Methods

#### Data Sets and Phylogenetic Analyses

A systematic search for clock components was conducted, building on previous studies exploring the evolution, duplication, and loss of circadian clock components (Thakkar et al. 2022; Kotwica-Rolinska et al. 2022). To identify circadian clock proteins and genes encoding dTIM, mTIM, TOF1, PERIOD, dCRY, mCRY, 6-4 Photolyase, JETLAG, and BRWD3 in Bilateria/Metazoa, GenBank (NCBI) protein and genomic databases, as well as transcriptome shotgun assemblies (TSA), were utilized. BLASTP and tBLASTn algorithms were applied with taxon-restricted searches targeting specific lineages at the levels of orders, suborders, infraorders, and, in some cases, families. In certain instances, genome or whole-genome shotgun contigs (wgs) were also examined. Protein sequences of circadian clock genes from *Drosophila melanogaster*, *Danaus plexippus*, and *Pyrrhocoris apterus* served as initial queries. Reciprocal and lineage-focused searches incorporated queries representing identified proteins from related taxa. To detect duplicate hits (common in TSAs) or closely related proteins (e.g., bHLH-PAS proteins instead of PERIOD), the E-INS-i algorithm in MAFFT was used for alignment, followed by FastTree analysis (Price et al. 2009), both conducted in Geneious Prime 21.0.3 (Biomatters, New Zealand).

### Phylogenetic analyses

To identify specific types of TIM, CRY, FBXL, or PAS proteins (e.g., distinguishing PER from bHLH PAS proteins), sequences were aligned using the MAFFT algorithm in Geneious Prime 21.0.3 (Biomatters, New Zealand). Representative datasets containing target proteins and related types were included. Ambiguously aligned regions were trimmed, and phylogenetic analyses were conducted using RAxML with a maximum likelihood GAMMA-based model in Geneious Prime 21.0.3.

The Metazoan phylogenies presented in Figs. 2, 4, 5, 6, 7, 8 and Fig. S8 were retrieved using TIMETREE 5 (https://timetree.org/; Kumar et al. 2022) and cross-referenced with recent molecular phylogenomic studies. These included works focused on insects (Misof et al. 2014), chelicerates, and crustaceans (von Reumont et al. 2012; Thomas et al. 2020; Bernot et al., 2023). For insects, phylogenomic studies specific to Polyneoptera (Wipfler et al. 2019), the hemipteroid assembly (Johnson et al. 2018), and Coleoptera (McKenna et al. 2019) were used to refine the corresponding sections of the phylogeny.

### Gene Loss

While it is impossible to definitively prove the absence of a gene, in some cases, gene loss is the most plausible explanation. Recent advances in phylogenomics, the availability of extensive TSA data, and an increasing number of sequenced genomes have enabled a systematic exploration of circadian clock genes across major Bilateria groups (Protostomia and Deuterostomia). Our analysis focuses on lineage-specific gene losses that are strongly supported by data from multiple species, whole-genome assemblies, and deep transcriptome sequencing. Evidence for gene loss is summarized for specific genes and animal groups in Supplementary Table 1.

#### Prediction of Protein Domains

Domains of the *Drosophila melanogaster* dTIM protein were annotated based on boundaries defined by Lin et al. (2023) and cell-based experiments by Saez and Young (1996). These domains include the dCRY-interaction domains, ARMADILLO 1 (ARM1), ARMADILLO 2 (ARM2), PER-binding sites 1 and 2 (PER-bind #1, PER-bind #2), as annotated in the *D. melanogaster* dTIM isoform P (NP_001334730).

All dTIM sequences in our dataset were aligned with *D. melanogaster* dTIM using MAFFT to identify regions of similarity. In fig. 2, these similarities are visualized as color intensities corresponding to numerical values. Supplementary figures include detailed protein models with specific similarity values next to annotated domains. In gene models (Figure 6 and 7), only two shades are used: regions with >45% similarity for ARM1, ARM2, and PER-binding sites are highlighted, while 38% similarity in *Limulus* is labeled as "low similarity" with a paler shade. The highly variable CRY-interaction domain at the C-terminal tail is highlighted for similarities >25%, with values as low as 19% in *Bemisia* indicated as "low similarity."

Nuclear localization signal (NLS) domains were predicted for each protein sequence using psort II (https://psort.hgc.jp/) and NLStradamus (http://www.moseslab.csb.utoronto.ca/NLStradamus/) prediction software. Acidic domains were annotated based on the following criteria: the sequence must be at least half the size of the reference sequence, located in the corresponding protein region, contain at least 20% acidic residues (D, E), and fewer than 10% basic residues (K, H, R). To further investigate acidic domains, sequence motifs were identified using Gapped Local Alignment of Motifs (GLAM2) within the MEME Suite (https://meme-suite.org/meme/). Two regions of 35 insect TIM proteins were scanned: sequences between armadillo domains #1 and #2, and the C-terminal tails.

TIM protein sequences analyzed were identical to those in Fig. S2. dTIM sequences were aligned using MAFFT (Katoh and Stanley 2013) (v7.490, Algorithm: E-INS-i, scoring matrix: BLOSUM62) in Geneious Prime to define the regions for analysis. The region between armadillo domains corresponds to amino acids E249-T577 in *Drosophila melanogaster* TIM and includes the putative PER-binding site #1. Insect dTIM C-tails were defined as the amino acids following residues K1117 and F1118 in *D. melanogaster* dTIM (isoform P served as the reference). These residues are highly conserved across most insect dTIM proteins, and the C-tail region includes the entire putative CRY-binding domain defined in *D. melanogaster*. The C-tail has been lost or reduced in some insect taxa. Two conserved motifs (~30 amino acids and ~15 amino acids) were identified in 33 and 24 insect species, respectively, and are presented as alignments (MAFFT) or alignments with 12–13 flanking amino acids in Supplementary Fig. S4. Amino acid polarity is marked by distinct color coding, with acidic residues in the conserved motif further highlighted by black borderlines.

To calculate the length of the variable region in the central part of the protein and the C-terminal variable region (both shown in Fig. 2), we aligned all protein sequences in our dataset using MAFFT in order to identify conserved motifs. The following conserved motifs were found near the analyzed regions: YKDQ, located at the end of the dCRY-interaction domain; LLLR, marking the start of PER-binding site #2; and DLIE, situated in the latter half of the PER-binding site #2 sequence. Distances between these specific sites were measured using Geneious Prime 2024 software (Biomatters, Auckland, New Zealand). The values for each sequence distance were plotted as individual dots in Prism 7 (GraphPad Software, La Jolla, CA, USA) for all proteins in the dataset.

#### Substitutions per Amino Acid per Million Years

To calculate substitution rates per amino acid position for dTIM and PER proteins, we employed the following approach. Protein sequences (dTIM or PER) were aligned using the E-INS-i algorithm in MAFFT (Katoh and Standley 2013). The complete alignments were used to infer phylogenetic trees with RAxML (Stamatakis 2014) under the PROTGAMMAJTT model, ensuring the topology matched the evolutionary relationships of the organisms. Constraint trees were created in

TreeGraph 2 (Stöver and Müller 2010). For Crustacea, where phylogenies are still debated, we enforced monophyly with insects but did not specify internal branching. Similarly, Polyneoptera were constrained as monophyletic without defining their internal topologies.

The resulting unrooted trees were swapped to position Protostomia and Deuterostomia as sister groups. Branch lengths were extracted and summed from the Protostomia/Deuterostomia split to terminal species. These values were then divided by 700 million years (the estimated divergence time of Protostomia and Deuterostomia) to compute substitution rates per amino acid position per million years. The values were plotted in Prism 7 (GraphPad Software, La Jolla, CA, USA), with each dot on the plot representing a single protein.

#### Gene Models

In brief, similarities in gene structure were assessed as previously (Smykal et al. 2020) when the gene models were either directly downloaded from GenBank species' Whole-genome shotgun contigs (wgs) or Representative genomes (RefSeq genomes). Alternatively, gene models were manually reconstructed using genomic and transcriptomic data from GenBank, following two distinct scenarios: (1) Long genomic contigs without annotated *timeless* genes. The wgs contigs were manually annotated by mapping Transcriptome Shotgun Assembly (TSA) and/or non-redundant nucleotide (nr/nt) *d-tim* sequences by the Minimap2 mapper (Li 2021) within Geneious Prime 2024 software (Biomatters, Auckland, New Zealand). *Lethocerus indicus tim* gene model was reconstructed by mapping TSA sequences from three related species: *Trichocorixa calva, Gelastocoris ocwulatus, Buenoa margaritacea*, to an unannotated genomic scaffold. Similarly, *Neoneuromus ignobilis tim* gene was annotated using TSA from *Protohermes xanthodes*. In *Pogonus chalceus*, two partial non-overlapping TSA sequences cover the majority of the coding sequence (CDS). The missing region spanning parts of exons 7 and 8 was reconstructed by mapping TSAs from related beetle species *Amphizoa insolens, Tribolium castaneum, and Sinaspidytes wrasei*. (2) Fragmented genomic contigs without annotated *timeless* genes. The *tim* exons were annotated and contigs scaffolded using *tim* CDS or TSA sequences. In the case of *Lepisma saccharina*, *Thermobia domestica d-tim* TSAs were mapped to the seven *L. saccharina* unannotated contigs. See Table S5 for the accession number of genomic and TSA sequences used for gene reconstruction and Table S6 for accession numbers used to annotate the CDS in gene models.

#### Exon Homology, Similarity, and dTIM Domain Localization in Gene Models

The *tim* CDSs of representative species were translated and aligned using MAFFT (Katoh and Standley 2013) within Geneious Prime 2024 software to determine conserved domains and conserved/homologous exons and exon boundaries. Two protein alignments were used: "species-to-species pair alignments" and "all representative proteins alignment". The longest *d-tim* isoforms of 29 representative species were analyzed in figure 6. Available *d-tim* isoforms were included in figure 7, where 28 TIM sequences from 11 dipteran species were aligned.

Two exon boundaries were determined as conserved through the species-to-species dTIM/dTIM pair alignment if a minimum of three out of five amino acids (aa) located at the exon/intron boundary were identical. The similarity of amino acids was analyzed within: One pair of homologous exons if at least one side of the exon border could be aligned, or several exons homologous to one 'fused' exon if both sides of the exon border could be aligned, according to species-to-species pair exon alignment using MAFFT (% Similarity, auto algorithm, scoring matrix: BLOSUM90 with threshold 1). A semi-quantitative scale with five similarity ranges (30-40; 41-50, 51-60, 61-70; 71-100 %) was used to visualize exon/exon similarity as quadrilaterals of five different levels of greyscale in figures 6 and 75.

CRYPTOCHROME (CRY)-interaction domains and PERIOD (PER)-binding sites were mapped and annotated in dTIM protein representative sequences utilizing domains defined in *Drosophila melanogaster* dTIM (Lin et al. 2023; Wulbeck et al. 2005). The figure was drawn using Adobe Illustrator (version 6) software.

#### Gene Syntenies

In brief, gene syntenies were performed as was done in a recent study exploring gene duplications (Smykal and Dolezel 2023). To compare gene synteny of genes neighboring *Drosophila*-type *cryptochrome* (*d-cry*) in heteropteran species *Apolygus lucorum*, *Lygus lineolaris* and *Lethocerus indicus* with species lacking *d-cry*, i.e. *Cimex hemipterus* and *Riptortus pedestris*, we identified protein-coding genes surrounding *d-cry* in the annotated *A. lucorum*, and non-annotated *L. indicus* genomes, and then search for those genes in the genomes of species lacking *d-cry*.

*L. hesperus d-cry*-encoding Transcribed Sequence Assembly (TSA, acc # GDHC01010508.1) was used to identify and map *d-cry* gene in *A. lucorum* annotated genomic contig by BLAST (Basic Local Alignment Search Tool). TSAs of two protein-coding genes upstream of *d-cry*: *Axin* (*Axn*, GE061_006998) and *yippee-like* (*yippee*, GE061_006997), and two downstream genes: *DDRGK domain-containing protein 1* (*Ddrgk1*, GE061_006990) and *eclair* (*eca*, GE061_006989) were searched in *Ly. hesperus* and *Ly. lineolaris* TSA databases using *A. lucorum* queries to identify and map corresponding genes in non-annotated *L. lineolaris* genomic contig and verify (and correct) original *A. lucorum* genes. *Axn*, *yippee*, *Ddrgk1,* and *eca* genes were searched and identified in *L. indicus*, *C. hemipterus* and *Ri. pedestris* TSA databases. TSAs were used to map the genes in their genomic contigs.

To identify the *d-cry* locus in the non-annotated *L. indicus* genome, the *Le. indicus* genome was searched by BLAST using the *Ly. hesperus* d-CRY protein sequence. Next, *L. indicus d-cry* contig (JACDQD010000008.1) was (pre-)annotated using the plugin Augustus (version 0.1.1) within Geneious Prime (2024.0.5) with *Rhodnius prolixus* as a reference species. *d-cry* and *d-cry* upstream: *venom protease-like* (*vp-l*), *WD repeat-containing protein 19* (*wdr19*), *Ubiquitin carboxyl-terminal hydrolase 7* (*Usp7*) and *inositol polyphosphate 5-phosphatase E* (*inpp5e*) and d-cry downstream: *Serine hydroxymethyltransferase* (*Shmt*) predicted genes' coding sequences (CDS) were used as queries for searching for *Le. indicus* TSAs, which were mapped back to the contig to correct the predicted gene models. CDSs of corrected upstream and downstream genes were searched in *Lygus*, *Cimex* and *R. pedestris* TSA databases and mapped to *y. lineolaris*, *A. lucorum* (using *Lygus* TSAs), *C. hemipterus,* and *R. pedestris* genomic contigs.

The same approach applied to *d-cry* was used for *jetlag* (*jet*) gene synteny, including the same species set used for TSA and genomic search and gene mapping, respectively. *L. hesperus jet* TSA (acc # GBRD01002318) was used to identify and map *jet* gene in *A. lucorum* annotated genomic contig using BLAST. TSAs of three protein-coding genes upstream of *jet*: *histone acetyltransferase kat2a* (*kat2a*, GE061_007944), *transmembrane protein 59-like* (*tmem59*, GE061_007947), *ATP-dependent (S)-NAD(P)H-hydrate dehydratase* (*Naxd*, GE061_007948) and one downstream gene: *Dual specificity protein phosphatase* (*Mkp3*, GE061_007950) were searched in *L. hesperus* and *L. lineolaris* TSA databases and mapped to *L. lineolaris* and *A. lucorum* genomic contigs.

*jetlag* gene synteny in *Le. indicus* was performed the same way as for *d-cry*. The *jetlag*-containing genomic contig (JACDQD010000012.1) was searched for surrounding protein-coding genes and four upstream genes: *serine/arginine repetitive matrix protein 1* (*srrm1*), *ribosomal RNA processing protein 1 homolog* (*rrp1*), *serine/threonine-protein phosphatase 4 regulatory subunit 4* (*ppp4r4*) and *tyrosine-protein kinase Shark* (*Shark*), and three downstream genes: *facilitated trehalose transporter Tret1*

(*Tret1*), *PIN2/TERF1-interacting telomerase inhibitor 1* (*pinx1*) and *Translational activator of cytochrome c oxidase 1* (*Taco1*), were searched in other four analyzed species.

Accession numbers of genomic contigs and representative TSAs are in Supplementary Tables S3 and 4. The final gene syntenies were drawn in CorelDRAW X6 (16.4.0.1280).

#### Table S1 *d-timeless* (*d-tim*), *period*, *cry*, and *jet* Gene Losses (August 2024)

TSA… transcriptome shotgun assemblies
wgs… whole-genome shotgun contigs

| **Taxonomy** | **Data suggesting *d-tim* loss as the most parsimonious explanation** |
|---|---|
| Hymenoptera (NCBI:txid7399) | Heavily sequenced group (both transcriptomes and genomes) with well-assembled genomes. No dTIM found (the only hits were mTIM sequences), >400 genomes |
| Isoptera (Termitoidea, termites) (NCBI:txid1912919) | Heavily sequenced group (both transcriptomes and ongoing genome projects) with well-assembled and compact genomes. With the exception of *Porotermes*, an early-branching termite species for which dTIM TSA hit was found (Kotwica-Rolinska et al. 2022; the hit branches within Blattodea dTIMs), no *d-tim* hit has been found in either transcriptomes or genomes of remaining available species (including *Zootermopsis nevadensis*). Thus, *d-tim* gene/ pseudogene might be present in some early termites but has been lost in the majority of species. 6 genomes at NCBI |
| Amphipoda (NCBI:txid6821) | No hit (even partial) in either TSA, genomes, or wgs. However, only circa 9 genomes have been sequenced so far for Amphipoda, furthermore, the genomes seem variable in size (Rees et al. 2007) including relatively large ones, such as 3.6 Gb for *Parhyale hawaiensis* (Kao et al. 2016) |
| Scorpiones (NCBI:txid6855) | No hit (even partial) in either TSA, genomes, or wgs. However, only circa 4 genomes have been sequenced so far for Scorpiones (*Centruroides sculpturatus* seem to be well annotated) |
| Chordata (NCBI:txid7711) | Heavily sequenced group (both transcriptomes and genomes) with well-assembled genomes. No dTIM hits (the only hits were mTIM sequences), >1000 genomes |
| Platyhelminthes (NCBI:txid6157) | Heavily sequenced group (both transcriptomes and genomes) with well-assembled genomes. No dTIM found (the only hits were mTIM sequences), >60 genomes |
| Nematoda (NCBI:txid6231) | Heavily sequenced group (both transcriptomes and genomes) with well-assembled genomes. No dTIM found (the only hits were mTIM sequences), >200 genomes |
| Cnidaria (NCBI:txid6073) | No hits (only mTIM), >100 genomes |
| Placozoa (NCBI:txid10226) | No hits (only mTIM), 2 genomes |
| Porifera (NCBI:txid6040) | No hits (only mTIM), >20 genomes |
| Green plants (NCBI:txid33090) | No hits (only TOF1), >1000 genomes |
| Diatoms (NCBI:txid2836) | No hits (only TOF1), >70 genomes |
| Fungi (NCBI:txid4751) | No hits (only TOF1, also known as Swi1 in *Schisosacharomyces pombe*), >5000 genomes available |
| **Taxonomy** | **Data suggesting *period* loss as the most parsimonious explanation** |
| Echinodermata (NCBI:txid7586) | No PER hits, >60 genomes |
| Hemichordata (NCBI:txid10219) | No PER hits, 2 genomes |

| **Taxonomy** | **Data suggesting *m-cry* loss as the most parsimonious explanation** |
|---|---|
| Subset of Diptera families:<br>Drosophilidae (NCBI:txid7214),<br>Muscidae (NCBI:txid7366),<br>Tephritidae (NCBI:txid7211),<br>Diopsidae (NCBI:txid139644) | no mCRY hits (only 6_4 Photolyase hits)<br>Well-sequenced and annotated group:<br>Drosophilidae >300 genomes,<br>Muscidae >10 genomes,<br>Tephritidae >25 genomes,<br>Diopsidae 3 genomes |
| **Taxonomy** | **Data suggesting *jetlag* loss as the most parsimonious explanation** |
| Subset of Coleoptera, following infraorders:<br>Cucujiformia (NCBI:txid41088)<br>Scarabaeiformia (NCBI:txid41086)<br>Staphyliniformia (NCBI:txid41085) | No hit in either TSAs or genomes<br><br>Cucujiformia: 136 genomes<br>Scarabaeiformia: 36 genomes<br>Staphyliniformia> 42 genomes |
| Hymenoptera (NCBI:txid7399) | Heavily sequenced group (both transcriptomes and genomes) with well-assembled genomes. No JET found; >400 genomes |
| Psocodea (NCBI:txid1930602) | No hit in either TSA or genomes (12 available) |
| Pentatomomorpha (NCBI:txid33357) | No hit in either TSA or genomes (13 available) |
| Subset of Cimicomorpha, families:<br>Cimicidae (NCBI:txid30078)<br>Reduvidae (NCBI:txid27479) | No hit in either TSAs or genomes<br>Cimicidae: 2 genomes<br>Reduvidae: 2 genomes |
| Aphidomorpha (NCBI:txid33380) | No hit (even partial) in either TSAs, genomes, or wgs.<br>38 genomes available |
| Amphipoda (NCBI:txid6821) | No hit (even partial) in either TSAs (transcriptome shotgun assemblies), genomes, or wgs. However, only circa 9 genomes have been sequenced so far for Amphipoda, furthermore, the genomes seem variable in size (Rees et al. 2007) including relatively large ones, such as 3.6 Gb for *Parhyale hawaiensis* (Kao et al. 2016) |
| Parasitiformes (NCBI:txid6934)<br>A subset of Acari including ticks | No hit in either TSAs or genomes: 31 genomes available |
| **Taxonomy** | **Data suggesting *d-cry* loss as the most parsimonious explanation** |
| Subset of Coleoptera, following infraorders:<br>Cucujiformia (NCBI:txid41088)<br>Scarabaeiformia (NCBI:txid41086)<br>Staphyliniformia (NCBI:txid41085) | No hit in either TSAs or in genomes<br><br>Cucujiformia: 136 genomes<br>Scarabaeiformia: 36 genomes<br>Staphyliniformia: 42 genomes |
| Phthiraptera (NCBI:txid85819), subset of Psocodea | No hit in either TSAs or genomes (8 available) |
| Pentatomomorpha (NCBI:txid33357) | No hit in either TSAs or genomes (13 available) |
| Subset of Cimicomorpha, families:<br>Cimicidae (NCBI:txid30078)<br>Reduvidae (NCBI:txid27479) | No hit in either TSAs or genomes<br><br>Cimicidae 2 genomes<br>Reduvidae 2 genomes |

| | |
|---|---|
| Amphipoda<br>(NCBI:txid6821) | No hit (even partial) in either TSAs (transcriptome shotgun assemblies), genomes, or wgs. However, only circa 9 genomes have been sequenced so far for Amphipoda, furthermore, the genomes seem variable in size (Rees et al. 2007) including relatively large ones, such as 3.6 Gb for *Parhyale hawaiensis* (Kao et al. 2016) |
| Scorpiones<br>(NCBI:txid6855) | No hit (even partial) in either TSAs (transcriptome shotgun assemblies), genomes, or wgs. However, only circa 4 genomes have been sequenced so far for Scorpiones (*Centruroides sculpturatus* seem to be well annotated) |
| Parasitiformes (NCBI:txid6934)<br>A subset of Acari including ticks | No hit in either TSAs or genomes:<br>31 genomes available |
| Chordata<br>(NCBI:txid7711) | Heavily sequenced group (both transcriptomes and genomes) with well-assembled genomes. No dCRY hits (the only hits were mCRY sequences), >1000 genomes |

#### Table S2. dTIM Gene Duplications

| **Taxonomy** | **Species** | **DNA accession number; Gene ID** | **Transcript accession number** | **Note** |
|---|---|---|---|---|
| Crustacea; Multicrustacea; Copepoda | *Lepeophtheirus salmonis* | LOC121131333 | XM_040726811.1 | model in Supplementary fig. 5 |
| | | LOC121130538 | XM_040726275.1 | model in Supplementary fig. 5 |
| | | LOC121130369 | XM_040726120.1 | model in Supplementary fig. 5 |
| Crustacea; Branchiopoda | *Daphnia magna* | LOC116920145 | XM_045172064.1 | model in Supplementary fig. 6 |
| | | LOC116923752 | XM_045178858.1 | model in Supplementary fig. 6 |
| | | LOC116916080 | XM_032921270.2 | model in Supplementary fig. 6 |
| | | LOC116915790 | XM_045168213.1 | model in Supplementary fig. 6 |
| Chelicerata | *Limulus polyphemus* | LOC106464343 | XM_022392591.1 | model in Supplementary fig. 4 |
| | | LOC106469873 | XM_022398446.1 | model in Supplementary fig. 4 |

#### Table S3. Synteny in the *d-cry* Locus (Depicted in Fig. 3G and S12A)

| **species + contig acc #** | **Gene or cDNA/TSA, species, acc #** |
|---|---|
| *Lygus lineolaris JAEMON010000012.1* | *d-cry Lygus hesperus* GDHC01010508.1 |
| | *Ddrgk1 Lygus hesperus* GBHO01041210.1 |
| | *eca Lygus hesperus* GBHO01005011.1 |
| | *inpp5e Lygus hesperus* GBHO01009246.1 |
| | *Axn Lygus hesperus* GDHC01015611.1 |
| | *Shmt Lygus lineolaris* GCXM01032240.1 |
| | *Usp7 Lygus hesperus* GDHC01019830.1 |
| | *wdr19 Lygus hesperus* GDHC01020915.1, GBHO01000933.1 |
| | *yippee Lygus hesperus* GBHO01041662.1 |
| Apolygus lucorum WIXP02000015.1 | *d-cry Lygus hesperus* GDHC01010508.1 |
| | *Ddrgk1 Lygus hesperus* GBHO01041210.1 |
| | *eca Lygus hesperus* GBHO01005011.1 |
| | *inpp5e Lygus hesperus* GBHO01009246.1 |
| | *Axn Lygus hesperus* GDHC01015611.1 |
| | *Shmt Lygus lineolaris* GCXM01032240.1 |
| | *Usp7 Lygus hesperus* GDHC01019830.1 |
| | *wdr19 Lygus hesperus* GDHC01020915.1, GBHO01000933.1 |
| | *yippee Lygus hesperus* GBHO01041662.1 |
| *Cimex hemipterus* | *d-cry – not found* |
| *Cimex hemipterus* JASJUQ010001410 | *Ddrgk1 Cimex lectularius* GBYH01012153.1 |
| | *eca Cimex lectularius* GBYH01012146.1 |
| | *inpp5e Cimex lectularius* GBYH01024336.1 |
| | *Axn Cimex lectularius* GBYH01036032.1 |
| | *Shmt Cimex lectularius* GDEK01028996.1 |
| | *Usp7 Cimex lectularius* GDEK01029216.1 |
| | *wdr19 Cimex lectularius* GBYH01036066.1 |
| | *yippee Cimex_lectularius* GDEK01001986.1 |
| | *vp-like Cimex lectularius* GDEK01004837.1 |
| *Lethocerus indicus* JACDQD010000008.1 | *d-cry Lethocerus indicus* GJWB01026588.1 |
| | *inpp5e* Lethocerus indicus GJWB01006126.1, GJWB01002010.1, GJWB01007124.1, partial coverage |
| | *Axn Lethocerus indicus GJWB01027068.1* |
| | *Shmt Lethocerus indicus* GJWB01023078.1, GJWB01016931.1 |
| | *Usp7 Lethocerus indicus* GJWB01030811.1 |
| | *wdr19 Lethocerus indicus* GJWB01007328.1, *Belostoma flumineum* GCWK01032557.1, *Lethocerus indicus* GJWB01013208.1, *Belostoma flumineum* GCWK01029820.1, *Lethocerus indicus* GJWB01007503.1, *Lethocerus indicus* GJWB01014013.1, *Belostoma flumineum* GCWK01023851.1, partial coverage |
| | *yippee Lethocerus indicus* GJWB01014554.1, GJWB01017227.1 |
| | *vp-like Lethocerus indicus* GJWB01025284.1 |
| | *vp-like Lethocerus indicus* GJWB01023677.1 |
| *Lethocerus indicus* JACDQD010000004.1 | *Ddrgk1 Lethocerus indicus* GJWB01016661.1 |
| | *eca Lethocerus indicus GJWB01025778.1* |
| *Riptortus pedestris* | *d-cry – not found* |
| *Riptortus pedestris* JADPXZ010000003.1 | *inpp5e Riptortus pedestris* GKOW01017232.1 |
| | *Axn Riptortus pedestris* GKOW01000064.1 |
| | *Shmt Riptortus pedestris* GKOW01106970.1 |
| | *Usp7 Riptortus pedestris* GKOW01024907.1 |
| | *wdr19 Riptortus pedestris* GKOW01051062.1 |
| | *yippee Riptortus pedestris* IADR01001218.1 |
| | *vp-like Riptortus pedestris* IADR01005779.1 |
| | *vp-like Riptortus pedestris* GKOW01028604.1 |
| *Riptortus pedestris* JADPXZ010000006.1 | *Ddrgk1 Riptortus pedestris* GKOW01047532.1 |
| | *eca Riptortus pedestris* IADR01006181.1 |

#### Table S4. Synteny in the *jetlag* Locus (Depicted in Figs. 3H and S12B)

| **species + contig acc #** | **Gene or cDNA/TSA, species, acc #** |
|---|---|
| *Lygus_lineolaris_-JAEMON010000014.1_annotated* | *jet Lygus hesperus GBRD01002318* |
| | *kat2a Lygus hesperus* GDHC01018225.1 |
| | *tmem59 Lygus hesperus* GDHC01002915.1 |
| | *Naxd Lygus hesperus* GBHO01007829.1 |
| | *Mkp3 Lygus hesperus* GBHO01007408.1 |
| *Lygus lineolaris - JAEMON010000009.1* | *srrm1 Lygus hesperus* GBRD01003465.1 |
| | *ppp4r4 Lygus hesperus* GBHO01041230.1 |
| | *rrp1 Lygus hesperus* GBHO01023925.1 |
| Lygus lineolaris - JAEMON010000007.1 | *Shark Lygus hesperus* GBHO01012115.1 |
| | *Tret1 Lygus hesperus* GBHO01016262.1 |
| | *pinx1 Lygus hesperus* GDHC01003347.1 |
| | *Taco1 Lygus hesperus* GBHO01011356.1 |
| Apolygus lucorum - WIXP02000016.1 | *jet Lygus hesperus GBRD01002318* |
| | *kat2a Lygus hesperus* GDHC01018225.1 |
| | *tmem59 Lygus hesperus* GDHC01002915.1 |
| | *Naxd Lygus hesperus* GBHO01007829.1 |
| | *Mkp3 Lygus hesperus* GBHO01007408.1 |
| *Apolygus lucorum - WIXP02000007.1* | *srrm1 Lygus hesperus* GBRD01003465.1 |
| | *ppp4r4 Lygus hesperus* GBHO01041230.1 |
| | *rrp1 Lygus hesperus* GBHO01023925.1 |
| Apolygus lucorum - WIXP02000005.1 | *Shark Lygus hesperus* GBHO01012115.1 |
| | *Tret1 Lygus hesperus* GBHO01016262.1 |
| | *pinx1 Lygus hesperus* GDHC01003347.1 |
| | *Taco1 Lygus hesperus* GBHO01011356.1 |
| *Cimex hemipterus* | *jet* – not found |
| *Cimex hemipterus* JASJUQ010001630.1 | *kat2a Cimex lectularius* GDEK01033364.1 |
| | *tmem59 Cimex lectularius* GDEK01002463.1 |
| | *Mkp3 Cimex lectularius* GDEK01002822.1 |
| *Cimex hemipterus* JASJUQ010000882 | *Naxd Cimex lectularius* GDEK01006328.1 |
| | *srrm1 Cimex lectularius* GBYH01020661.1 |
| | *ppp4r4 Cimex lectularius* GDEK01006107.1 |
| | *rrp1 Cimex lectularius* GBYH01026227.1 |
| | *Taco1 Cimex lectularius* GDEK01001076.1 |
| | *Tret1 Cimex lectularius* GDEK01004770.1 |
| | *pinx1 Cimex lectularius* GDEK01002581.1 |
| *Cimex hemipterus* JASJUQ010001765.1 | *Shark Cimex lectularius* GBYH01016742.1, GDEK01004540.1 |
| *Lethocerus indicus* JACDQD010000012.1 | *jet – Lethocerus indicus* GJWB01027467 |
| | *srrm1 Lethocerus indicus* GJWB01027207.1 |
| | *ppp4r4 Lethocerus indicus* GJWB01029598.1, GJWB01011308.1 |
| | *rrp1 Lethocerus indicus* GJWB01027713.1 |
| | *Shark Lethocerus indicus* GJWB01022002.1, GJWB01024613.1 |
| | *Tret1 Lethocerus indicus* GJWB01023355.1 |
| | *pinx1 Lethocerus indicus* GJWB01026351.1 |
| | *Taco1 Lethocerus indicus* GJWB01023842.1 |
| *Lethocerus indicus* JACDQD010000001.1 | *kat2a Lethocerus indicus* GJWB01025288.1 |
| | *tmem59 Lethocerus indicus* GJWB01027540.1 |
| | *Mkp3 Lethocerus indicus* GJWB01007119.1, GJWB01007308.1, *Ilyocoris cimicoides* GDEW01021870.1, *Corixa punctata* GDDR01015381.1, part. coverage |
| *Lethocerus indicus* JACDQD010000004.1 | *Naxd Lethocerus indicus* GJWB01022366.1 |
| *Riptortus pedestris* | *jet – not found* |
| *Riptortus pedestris* JADPXZ010000005.1 | *Shark Riptortus pedestris* GKOW01014958.1 |
| | *Tret1 Riptortus pedestris* GKOW01060243.1 |
| | *pinx1 Riptortus pedestris* GKOW01106563.1 |
| | *Taco1 Riptortus pedestris* GKOW01063983.1 |

| | |
|---|---|
| *Riptortus pedestris* JADPXZ010000006.1 | *kat2a Riptortus pedestris* GKOW01120792.1 |
| | *tmem59 Riptortus pedestris* IADR01020554.1 |
| | *Mkp3 Riptortus pedestris* IADR01014866.1 |
| | *srrm1 Riptortus pedestris* GKOW01040265.1 |
| | *ppp4r4 Riptortus pedestris* GKOW01032603.1 |
| | *rrp1 Riptortus pedestris* GKOW01048183.1 |
| *Riptortus pedestris* JADPXZ010000002.1 | *Naxd Riptortus pedestris* IADR01006076.1 |

#### Table S5. Gene model reconstruction table

| | |
|---|---|
| **Gene model for *Lepisma saccharina*** | |
| Genomic DNA from *Lepisma saccharina* (Genome assembly AU_Lsac_1.0):<br>JAGDQP010102583 (7959 bp)<br>JAGDQP010014933 (43939 bp)<br>JAGDQP010335617 (2660 bp)<br>JAGDQP010139949 (6139 bp)<br>JAGDQP010171045 (5169 bp)<br>JAGDQP010165524 (5324 bp) | cDNA sequences from *Thermobia domestica*:<br>GASN02066370<br>GASN02066371<br>AB644410 |
| **Gene model for *Lethocerus indicus*** | |
| Genomic DNA from *Lethocerus indicus*: JACDQD010000005 | cDNA sequence from *Trichocorixa calva*: GCYZ01029016 |
| | cDNA sequence from *Gelastocoris oculatus*: GCXI01032419 |
| | cDNA sequence from *Buenoa margaritacea*: GCXE01042166 |
| **Gene model for *Neoneuromus ignobilis*** | |
| Genomic DNA from *Neoneuromus ignobilis*: JAMXXB010000001 | cDNA sequence from *Protohermes xanthodes*: GCRQ01030715 |
| **Gene model for *Pogonus chalceus*** | |
| Genomic DNA from *Pogonus chalceus*: CM008233 | cDNA sequences from *Pogonus chalceus*:<br>JU426733<br>JU418895 |
| | cDNA sequence from *Amphizoa insolens*: GFUZ01001244 |
| | cDNA sequence from *Sinaspidytes wrasei*: GDNH01031115 |
| | cDNA sequence from *Tribolium castaneum*: NC_007420, NP_001106934.1 |

#### Table S6. Gene Model Similarities (Depicted in Figs. 4, 5, and Supplementary Figs)

| Taxonomy | Species | DNA accession number; Gene ID | Transcript accession number | Note |
|---|---|---|---|---|
| Diptera; Brachycera; Muscomorpha; Ephydroidea | *Drosophila melanogaster* | NT_033779.5 (3493986..3508119, complement); Gene ID: 33571 | NP_001334730.1 | model in figs. 4 and 5 |
| | *Drosophila pseudoobscura* | LOC4816795 | | model in fig. 5 |
| | *Drosophila willistoni* | LOC6642215 | | model in fig. 5 |
| | *Drosophila mojavensis* | LOC6577215 | | model in fig. 5 |
| | *Drosophila novamexicana* | LOC115760251 | | model in fig. 5 |
| | *Chymomyza costata* | ABW71828.1 | | model in fig. 5 |
| Diptera; Brachycera; Muscomorpha; Muscoidea | *Musca domestica* | LOC101894072 | | model in fig. 5 |
| Diptera; Brachycera; Muscomorpha; Tephritoidea | *Bactrocera dorsalis* | LOC105233653 | XM_011215785.4 | model in figs. 4 and 5 |
| Diptera; Brachycera; Stratiomyomorpha | *Hermetia illucens* | LOC119655307 | | model in fig. 5 |
| Diptera; Nematocera; Culicomorpha; Culicoidea | *Aedes aegypti* | LOC5567962 | XM_021847388.1 | model in figs. 4 and 5 |
| Diptera; Nematocera; Culicomorpha; Chironomoidea | *Clunio marinus* | JQ011272 CLUMA_CG006930 (CRK93394.1) | | model in fig. 5 |
| Lepidoptera | *Bombyx mori* | NC_051361.1_LOC733065 | XM_038021742.1 | model in fig. 4 |
| | *Pararge aegeria* | LOC120635901 | XM_039907115.1 | model in fig. 4 |
| Coleoptera, Adephaga | *Pogonus chalceus* | See table S5 | | model in fig. 4 |
| Coleoptera, Polyphaga | *Tribolium castaneum* | NC_036201_GeneID:657878 | XM_008194761.2 | model in fig. 4 |
| Megaloptera | *Neoneuromus ignobilis* | See table S5 | | model in fig. 4 |
| Neuroptera | *Chrysoperla carnea* | LOC123292921 | XM_044873638.1 | model in fig. 4 |
| Psocodea; Phthiraptera | *Pediculus humanus corporis* | NW_002987879_GeneID:8232513 | XM_002432376.1 | model in fig. 4 |
| Thysanoptera | *Thrips palmi* | LOC117639213 | XM_034374658.1 | model in fig. 4 |
| Hemiptera; Sternorrhyncha, Aleyroidea | *Bemisia tabaci* | LOC109035505 | XM_019049158.1 | model in fig. 4 |
| Hemiptera; Sternorrhyncha; Aphidimorpha | *Daktulosphaira vitifoliae* | LOC126898779 | XM_050673113.1 | model in fig. 4 |
| | *Acyrthosiphon pisum* | LOC100328917 | XM_029485187.1 | model in fig. 4 |
| Hemiptera; Heteroptera; Penatatomomorpha | *Pyrrhocoris apterus* | MW662134.1 | GI:2168441080 | model in fig. 4 |

| | | | | |
|---|---|---|---|---|
| Hemiptera; Heteroptera; Nepomorpha | *Lethocerus indicus* | See table S5 | | model in fig. 4 |
| Orthoptera | *Schistocerca serialis cubense* | LOC126457362 | XM_050093601.1 | model in fig. 4 |
| Odonata | *Ischnura elegans* | LOC124168471 | XM_046546734.1 | model in fig. 4 |
| Zygentoma | *Lepisma saccharina* | See table S5 | | model in fig. 4 |
| Hexapoda; Collembola | *Orchesella cincta* | LJIJ01000023.1_Ocin01_01260 | ODN05422.1 | model in fig. 4 |
| Crustacea; Multicrustacea; Copepoda | *Lepeophtheirus salmonis* | LOC121131333 | XM_040726811.1 | model in Supplementary fig. 5 |
| | | LOC121130538 | XM_040726275.1 | model in Supplementary fig. 5 |
| | | LOC121130369 | XM_040726120.1 | model in Supplementary fig. 5 |
| Crustacea; Multicrustacea; Decapoda | *Homarus americanus* | LOC121869648 | XM_042371176.1 | model in fig. 4 |
| Crustacea; Branchiopoda | *Daphnia magna* | LOC116920145 | XM_045172064.1 | model in Supplementary fig. 6 and in fig. 4 |
| | | LOC116923752 | XM_045178858.1 | model in Supplementary fig. 6 |
| | | LOC116916080 | XM_032921270.2 | model in Supplementary fig. 6 |
| | | LOC116915790 | XM_045168213.1 | model in Supplementary fig. 6 |
| Crustacea; Oligostraca | *Darwinula stevensoni* | Dst b1v02 scaf01583_DSTB1V02_LOCUS7617 | GI:2041332297 | model in fig. 4 |
| Chelicerata | *Limulus polyphemus* | LOC106464343 | XM_022392591.1 | model in Supplementary fig. 4 |
| | | LOC106469873 | XM_022398446.1 | model in Supplementary fig. 4 and in fig. 4 |
| Annelida | *Owenia fusiformis* | CAIIXF020000004_OFUS_LOCUS7441 | OFUSG13978.1 | model in fig. 4 |
| Mollusca | *Crassostrea gigas* | LOC105333471 | XM_034474211.1 | model in fig. 4 |
| Echinodermata; Echinozoa | *Strongylocentrotus purpuratus* | LOC587080 | XM_786834.5 | model in fig. 4 |
| Echinodermata; Asterozoa | *Acanthaster planci* | LOC110981676 | XM_022239459.1 | model in fig. 4 |

#### Supplementary References

Bernot, JP, Owen CL, Wolfe JM, Meland K, Olesen J, Crandall KA. 2023. Major Revisions in Pancrustacean Phylogeny and Evidence of Sensitivity to Taxon Sampling. Molecular Biology and Evolution 40(8).

Johnson KP, Dietrich CH, Friedrich F, Beutel RG, Wipfler B, Peters RS, Allen JM, Petersen M, Donath A, Walden KKO, et al. 2018. Phylogenomics and the evolution of hemipteroid insects. Proc Natl Acad Sci U S A 115:12775-12780.

Katoh K, Standley DM. 2013. MAFFT multiple sequence alignment software version 7: improvements in performance and usability. Molecular Biology and Evolution 30:772-780.

Kao D, Lai AG, Stamataki E, Rosic S, Konstantinides N, Jarvis E, Di Donfrancesco A, Pouchkina-Stancheva N, Semon M, Grillo M, et al. 2016. The genome of the crustacean Parhyale hawaiensis, a model for animal development, regeneration, immunity and lignocellulose digestion. Elife 5.

Kotwica-Rolinska J, Chodakova L, Smykal V, Damulewicz M, Provaznik J, Wu BC, Hejnikova M, Chvalova D, Dolezel D. 2022. Loss of Timeless Underlies an Evolutionary Transition within the Circadian Clock. Molecular Biology and Evolution 39.

Kumar S, Suleski M, Craig JM, Kasprowicz AE, Sanderford M, Li M, Stecher G, Hedges SB. 2022. TimeTree 5: An Expanded Resource for Species Divergence Times. Molecular Biology and Evolution 39.

Li H. 2021. New strategies to improve minimap2 alignment accuracy. Bioinformatics 37(23): 4572-4574.

Lin C, Feng S, DeOliveira CC, Crane BR. 2023. Cryptochrome-Timeless structure reveals circadian clock timing mechanisms. Nature 617:194-199.

McKenna DD, Shin S, Ahrens D, Balke M, Beza-Beza C, Clarke DJ, Donath A, Escalona HE, Friedrich F, Letsch H, et al. 2019. The evolution and genomic basis of beetle diversity. Proc Natl Acad Sci U S A 116:24729-24737.

Misof B, Liu S, Meusemann K, Peters RS, Donath A, Mayer C, Frandsen PB, Ware J, Flouri T, Beutel RG, et al. 2014. Phylogenomics resolves the timing and pattern of insect evolution. Science 346:763-767.

Price MN, Dehal PS, Arkin AP. 2009. FastTree: computing large minimum evolution trees with profiles instead of a distance matrix. Molecular Biology and Evolution 26:1641-1650.

Rees DJ, Dufresne F, Glemet H, Belzile C. 2007. Amphipod genome sizes: first estimates for Arctic species reveal genomic giants. Genome 50:151-158.

Saez L, Young MW. 1996. Regulation of nuclear entry of the Drosophila clock proteins period and timeless. Neuron 17:911-920.

Smýkal V, Pivarči M, Provazník J, Bazalová O, Jedlička P, Lukšan O, Horák A, Vaněčková H, Beneš V, Fiala I, et al. 2020. Complex evolution of insect insulin receptors and homologous decoy receptors, and functional significance of their multiplicity. Mol Biol Evol. 37(6):1775–1789.

Smykal V, Dolezel D. 2023. Evolution of proteins involved in the final steps of juvenile hormone

Stamatakis A. 2014. RAxML version 8: a tool for phylogenetic analysis and post-analysis of large phylogenies. Bioinformatics 30:1312–1313.

Stöver BC, Müller KF. 2010. TreeGraph 2: Combining and visualizing evidence from different phylogenetic analyses. BMC Bioinformatics 11:7

Thakkar N, Giesecke A, Bazalova O, Martinek J, Smykal V, Stanewsky R, Dolezel D. 2022. Evolution of casein kinase 1 and functional analysis of new doubletime mutants in Drosophila. Front Physiol 13:1062632.

Thomas GWC, Dohmen E, Hughes DST, Murali SC, Poelchau M, Glastad K, Anstead CA, Ayoub NA, Batterham P, Bellair M, et al. 2020. Gene content evolution in the arthropods. Genome Biology 21:15.

von Reumont BM, Jenner RA, Wills MA, Dell'ampio E, Pass G, Ebersberger I, Meyer B, Koenemann S, Iliffe TM, Stamatakis A, et al. 2012. Pancrustacean phylogeny in the light of new phylogenomic data: support for Remipedia as the possible sister group of Hexapoda. Molecular Biology and Evolution 29:1031-1045.

Wipfler B, Letsch H, Frandsen PB, Kapli P, Mayer C, Bartel D, Buckley TR, Donath A, Edgerly-Rooks JS, Fujita M, et al. 2019. Evolutionary history of Polyneoptera and its implications for our understanding of early winged insects. Proc Natl Acad Sci U S A.

Wulbeck C, Szabo G, Shafer OT, Helfrich-Forster C, Stanewsky R. 2005. The novel Drosophila tim(blind) mutation affects behavioral rhythms but not periodic eclosion. Genetics 169:751-766.
